# Rigidity transitions in a 3D active foam model of cell monolayers with frictional contact interactions

**DOI:** 10.1101/2025.04.17.649329

**Authors:** Jef Vangheel, Herman Ramon, Bart Smeets

**Affiliations:** KU Leuven, Dept. of Biosystems, Division of MeBioS, B-3001 Leuven, Belgium

## Abstract

In morphogenesis and disease, biological tissues may exhibit diverse mechanical properties due to their capacity to switch between fluid-like to solid-like states through the (un)jamming transition. Here, we introduce a novel foam model to investigate how active mechanical properties and cellular interactions govern this transition in active cell monolayers. This model explicitly represents 3D cell shapes and describes cell-cell interactions via discrete interacting surfaces. Simulations reveal that cell-cell adhesive tension promotes tissue fluidization in high adhesive tissues, where it mainly promotes cell deformability, while it induces solidification in the low adhesive regime, where it prevents cell-cell debonding. Moreover, we study the dynamic role of adhesive ligand turnover through an effective intercellular friction. Through simulated shear experiments, we find that intercellular friction strongly suppresses neighbor exchanges, but does not lead to solid-like tissue properties. We discuss the implications of our results for understanding the relationship between unjamming and partial epithelial-mesenchymal transition, highlighting how differences in adhesion dynamics and intercellular friction may reconcile conflicting observations in tissue mechanics and cancer metastasis.

## I. INTRODUCTION

Biological tissues are active, living materials with mechanical properties that can vary dramatically depending on their context and functional requirements. Unlike inert materials, tissues are composed of active cells that generate forces and constantly remodel their connections with their neighbors. By tuning cell properties, tissues may undergo phase transitions, shifting from a fluid-like to a solid-like state, or vice versa [1–3]. These (un)jamming phase transitions play an important role in phenomena such as embryonic morphogenesis [4–10] and wound healing [11, 12], and disease processes such as asthma [13] and cancer metastasis [14–17].

Our theoretical understanding of the jamming transition in biological tissues mainly stems from vertex and Voronoi models [18]. They have demonstrated how tissue fluidity is governed by interfacial tension [19] and active cell motility [20]. In these models, a critical shape index — a geometric parameter that is a measure of cell elongation — predicts the jamming transition. While initially limited to confluent tissues, later extensions of these models also accounted for the presence of intercellular pores [21, 22]. For example, in their ‘active foam’ model, Kim et al. demonstrated that cell-cell adhesion favors fluidization in confluent tissues, as predicted by vertex models, but conversely, promotes solidification in porous tissues [23]. Vertex models have also been extended to 3D, to explore the active properties of bulk 3D tissues [24–27], or to study the behavior of curved epithelial shells and sheets [28, 29].

However, vertex family models treat tissues as collections of cell-cell interfaces rather than interacting surfaces. This makes it difficult to represent the dynamic properties of cell-cell contacts with adherent junctions, such as cadherins. These regulate contact expansion by locally inhibiting myosin contractility, thereby reducing the cell-cell interfacial tension [30–32]. Moreover, their bond strength and turnover rate also control the dynamics of cell-cell separation and relative cell sliding, producing an effective viscous cell-cell friction at long timescales [33–35]. Recent alternative modeling approaches represent cells as discrete interacting surfaces [36], offering a more detailed description of cell-cell interaction mechanics, albeit at the cost of increased computational complexity. Coupled with viscous active shell models [37, 38], they enable parameterization using measurable mechanical properties of the actomyosin cortex. The cortex, approximated as viscous at long timescales [39–42], is then modeled as a curved, contractile, viscous shell encapsulating an incompressible cytoplasm [43, 44].

Here, we present a deformable cell model (DCM) of active cell monolayers that explicitly represents the 3D shape of individual cells and models cell-cell interactions through discrete, interacting surfaces [45]. This way, the DCM intrinsically captures cell shape and tissue structure, including porosity, as determined by cell mechanical properties [46]. In simulations of this model, we study cell dynamics, tissue structure and viscoelastic properties. Furthermore, we explore the jamming transition in an epithelial monolayer in function of adhesive tension, cell motility, intercellular friction and cell density. We show that increased cell-cell adhesive tension fluidizes at high adhesive tension, while promoting solidification at low adhesive tension. Moreover, we derive and validate theoretical scaling regimes that predict tissue fluidity based on cell-scale mechanical properties in both regimes. Finally, we examine the influence of intercellular friction on tissue-scale mechanical behavior. Although intercellular friction strongly suppresses neighbor exchanges, we find that its effect is distinct from the classical jamming transition.

## II RESULTS

### A. Computational model of foam-like cell mechanics with contact interactions

Our model represents the long time-scale viscous and contractile behavior of adherent cells based on physical properties of the cell’s surface, which lumps the actomyosin cortex and the membrane, Fig. 1(a). This surface is modeled as a thin, curved shell with thickness *t*_*c*_, free surface tension *γ* and viscosity *η*_*c*_, surrounding an incompressible cytoplasm, with internal pressure *P*_*b*_ [45] and volume 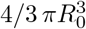. Cell-cell (*ω*_*cc*_) and cell-substrate (*ω*_*cs*_) adhesive tension represent the combined effect of adhesive ligands and the reduction of myosin contractility at the interface [30]. In our model, these effects are incorporated through i) a reduction of in-plane tension, thereby determining the contact angle, and ii) a normal adhesive traction, governing the mechanics of cell pull-off. Adhesive ligand turnover further introduces cell-cell (*ξ*_*cc*_) and cell-substrate (*ξ*_*cs*_) friction. At equilibrium, the balance of interfacial tension dictates individual cell shapes and monolayer architecture: Cell-cell adhesive tension increases the contact area with neighboring cells, while cell-substrate adhesive tension promotes substrate spreading. The shape of adherent cells changes from squamous to columnar cells with increasing *ω*_*cc*_*/γ* and decreasing *ω*_*cs*_*/γ*, Fig. 1(b). In an ideal hexagonal configuration, the equilibrium shape attained in simulations reproduces the theoretical aspect ratio obtained from tension balance when varying *ω*_*cs*_ and *ω*_*cc*_: 2*h/d* = (2*γ* − *ω*_*cs*_)*/*(*γ* − *ω*_*cc*_) (supplementary eq. S2), with height *h* and cell-cell distance *d*, Fig. 1(c).

**FIG. 1.**
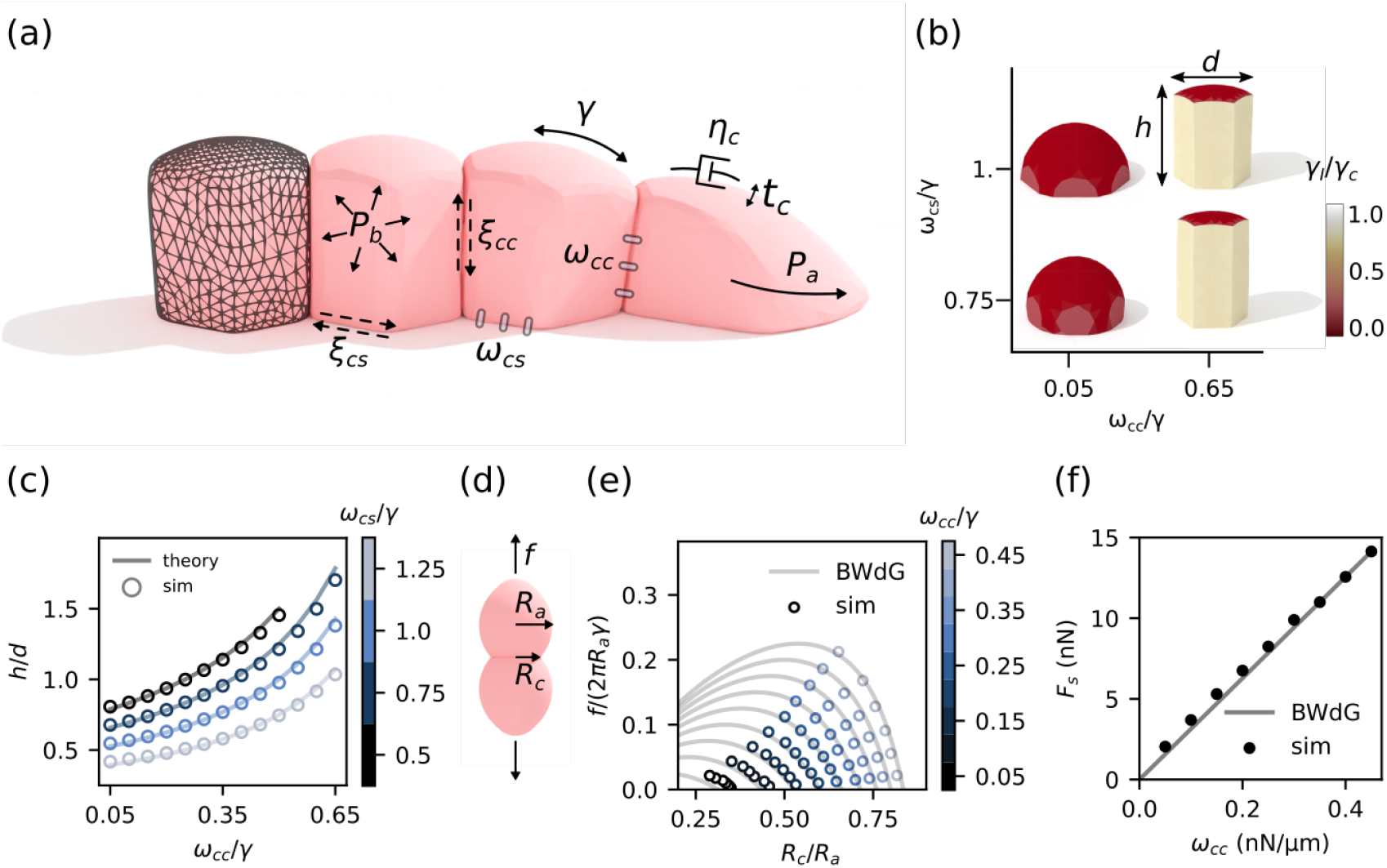
Overview of the computational deformable cell model. (a) DCM simulation of adhering cells on a substrate. Cell shape is outlined by a meshed surface (visualized on the left cell), which mechanically models the cell cortex with thickness *tc*, surface tension *γ* and viscosity *η*_*c*_, and cell volume is maintained through the internal pressure *P*_*b*_. Adhesive cell-cell (*cc*) and cell-substrate (*cs*) interactions are modeled by i) an adhesive contact pressure *ω* mechanically coupling the cells, ii) reduction of surface tension at the interface (*γ* − *ω*), and iii) an effective wet sliding friction *ξ*, accounting for dynamic bond turnover. Cell motility is modeled by an active pressure *P*_*a*_, eq. 5. (b) Simulated equilibrium cell shape ranges from squamous to columnar types by varying cell-cell (*ω*_*cc*_*/γ*) and cell-substrate adhesive tension (*ω*_*cs*_*/γ*). The color bar indicates local interfacial tension, *γ*_*I*_ = *γ* − *ω*_*ci*_. (c) Aspect ratio (cell height *h* over diameter *d*), of a simulated hexagonal cell plotted in function of *ω*_*cc*_*/γ* and *ω*_*cs*_*/γ*, and compared to the theoretical prediction 2*h/d* = (2*γ* − *ω*_*cs*_)*/*(*γ* − *ω*_*cc*_) (supplementary eq. S2). (d) Simulated dual micropipette aspiration of two adhering cells, and (e) comparison of simulated pull-off force in function of normalized contact radius *R*_*c*_*/R*_0_ to BwDG theory [47]. (f) Critical pull-off force in simulations follows the BwDG relation *F*_*s*_ = *πω*_*cc*_*R*_*a*_ ≈ *πω*_*cc*_*R*_0_ for adhesive vesicles.

Because our model explicitly captures interactions between distinct interfaces, it can accurately reproduce debonding and pull-off behavior during cell-cell separation — a key advantage that enables it to bridge the gap between classic adhesive sphere models and interface-based vertex models. By simulating a dual pipette aspiration experiment for cell-cell separation, varying relative cell-cell adhesive tension *ω*_*cc*_*/γ*, we show that our model replicates the theoretical prediction for adhesive vesicles as derived by Brochard-Wyart and de Gennes [47, 48], Fig. 1(e). Hence, our model reproduces the classical vesicle pull-off force *F*_*s*_ = *πω*_*cc*_*R*_0_, Fig. 1f.

### B. 3D structure and mechanical stress in static monolayers

We simulated cell monolayers by randomly initializing active cells on a substrate with periodic boundary conditions, followed by simulated annealing to achieve static configurations. We then characterized tissue structure and quantified the average hydrostatic tissue stress, *σ*_*h*_, based on intercellular forces using the virial theorem. Our 3D foam model is able to represent both confluent and (micro- and macro-)porous states by varying cell number density and adhesive tension *ω*_*cc*_, Fig. 2(a-b). At high number density, the tissue remains confluent (area density ≈ 1), with tissue stress transitioning from compressive to tensile as cell-cell adhesive tension increases, Fig. 2(a). At low number density, increasing cell-cell adhesive tension results in the nucleation and growth of intercellular pores, ultimately leading to phase separation into a locally confluent, arrested gel phase. In the absence of active (outward) traction forces, cell clusters within macro-porous tissue cannot sustain elevated tensile stress. Thus, tissue stress reaches its peak when the tissue is marginally confluent, coinciding with the onset of pore nucleation. Similarly, tissue stress increases with cell-substrate adhesive tension *ω*_*cs*_, Fig. 2(b), as area density increases with *ω*_*cs*_. At low cell-substrate adhesion and high cell-cell adhesive tension, the cell monolayer becomes mechanically unstable, leading to cell extrusion and multilayering, reminiscent of liquid film dewetting [49, 50]. It should be noted that, because our model restricts cell motility to the substrate, the tissue will not evolve toward mechanical equilibrium as described by the Young–Dupré equation. Rather, it stabilizes in an arrested gel-like state.

**FIG. 2.**
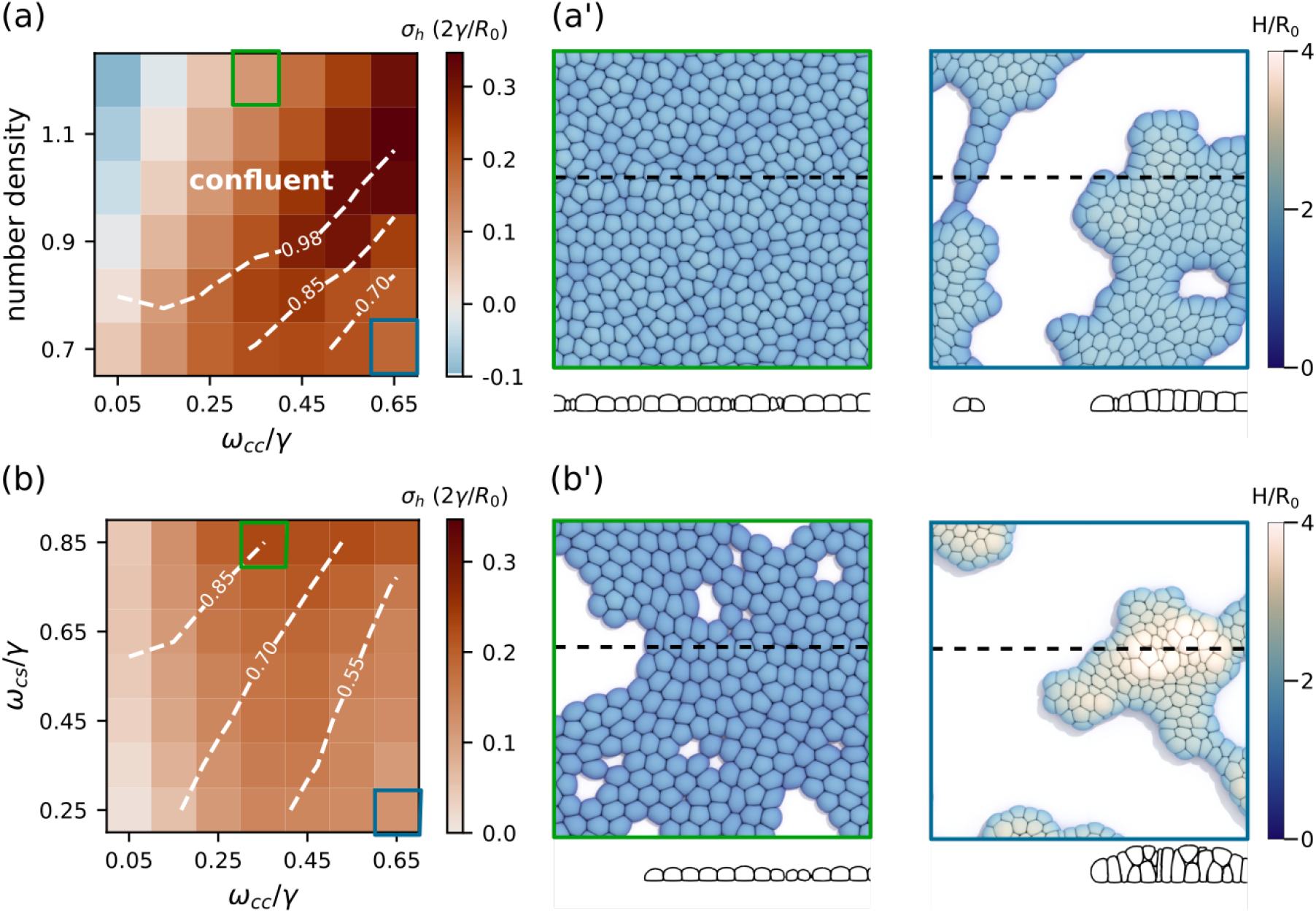
Structure and mechanical stress in quasi-static monolayers of 400 cells. (a) Normalized hydrostatic tissue stress *σ*_*h*_*R*_0_*/*2*γ* (positive values indicate tension, negative values indicate compression) for varying cell-cell adhesive tensions *ω*_*cc*_*/γ* and number densities at constant cell-substrate adhesive tension *ω*_*cs*_*/γ* = 0.825. Dashed white lines show iso-contours of 2D cell density. (a’) Visualizations of the apical and side view for parameters indicated by the blue and green box in (a). The dashed line indicates the position of the side view. The colorbar shows the relative tissue height *H/R*_0_. (b) Hydrostatic tissue stress for varying cell-cell and cell-substrate adhesive tensions at constant number density 0.7. Dashed white lines show iso-contours of 2D cell density. (b’) Visualizations of the apical and side view. Tissue dewetting can be observed at low *ω*_*cs*_*/γ* and high *ω*_*cc*_*/γ*.

### C. Cell deformation controls fluidity at high adhesive tension

Next, we quantified tissue fluidity in simulations of monolayers with actively migrating cells, undergoing rotational diffusion with persistence time *τ*_*p*_. The active force of cell migration may be parameterized by the dimensionless pressure *p*_eff_ = *p*_*a*_*R*_0_*/*(2*γλ*_*a*_), with active pressure *p*_*a*_ and dipole strength 1*/λ*_*a*_ (see methods). To characterize tissue fluidity in simulations, we measured 2D mean-squared relative displacements (MSRD) between initially neighboring cell pairs, Fig. 3a. By focusing solely on relative displacements, we eliminate the contribution of bulk tissue movements. These measures show that adhesive tension *ω*_*cc*_ can drive subdiffusive motion toward diffusive behavior, Fig. 3a. We quantified the effective diffusivity as *D*_eff_ = lim_*t*→∞_ MSRD(*t*)*/*4*t* (Fig. 3b), which decreases by orders of magnitude by decreasing motile pressure. However, *D*_eff_ changes non-monotonically with increasing *ω*_*cc*_, Fig. 3b. This is also reflected in cell trajectories, Fig. 3c, where cell movements become caged at intermediate values of cell-cell adhesive tension. Hence, we identify two distinct regimes: a low adhesive regime where *ω*_*cc*_*/γ* promotes solidification, and a high adhesive, confluent regime where *ω*_*cc*_*/γ* promotes fluidization.

**FIG. 3.**
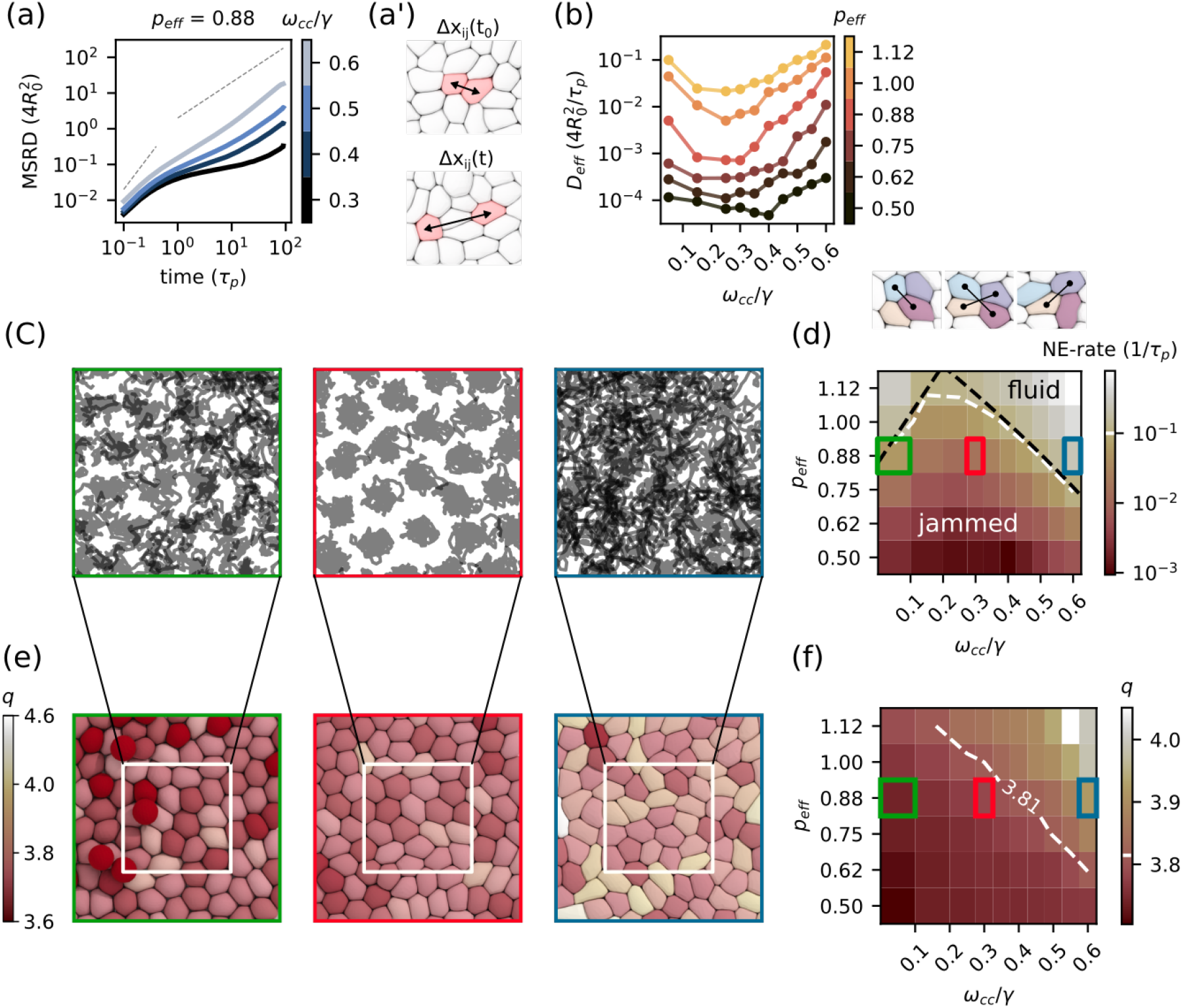
Fluidity of confluent tissue at number density 1.1. (a) Mean squared relative displacement (MSRD) between initially contacting cell pairs for varying *ω*_*cc*_*/γ*, with (a’) a visualization of the displacement between the cell pair *ij*. (b) Effective diffusion coefficient normalized by cell radius *R*_0_ and persistence time *τ*_*p*_. (c) Visualization of cell trajectories at *p*_eff_ = 0.875 and varying *ω*_*cc*_*/γ*. (d) Neighbor exchange rate (NE-rate) in the tissue as a measure for fluidity. The threshold that distinguishes solidfrom fluid-like tissue is set at NE-rate = 10^−1^(1*/τ*_*p*_), indicated by the white dashed line. The black dashed lines indicate scaling 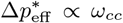 for the low adhesive regime, and 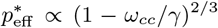 for the high adhesive regime. The inset shows an example of T1-transition in confluent tissue. (e) Visualization of the apical cell shape. Cells become more elongated by increasing cell motility and adhesive tension. See also supplementary Fig. S1 and supplementary video S1 for a visualization of the simulated tissues. (f) Apical cell shape in function of adhesive tension and cell motility for confluent tissue. The white line indicates the critical shape index *q*^∗^ = 3.81.

At high adhesive tension, we hypothesized that the ability of cells to elongate modulates the probability of T1-transitions and thereby regulates tissue fluidity. This follows predictions of vertex models, where fluidization occurs as the shape index, a measure of cell elongation, increases beyond a critical value. In our foam model, cell elongation may arise from both external (interactions with neighbor cells) and internal (active cell elongation and migration) forces. In both cases, the extent of elongation depends on the balance between (internally or externally generated) pressure and net cortex tension at the cell-cell interface, *γ*_eff_ = *γ* − *ω*_*cc*_. Restricting to internally generated active forces, we may estimate the 2D aspect ratio of a cell by equating the pressure difference over the surface at the long axis, including the active pressure dipole, and the short axis (derivation provided in the supplementary text). Assuming that the tissue unjams when a critical aspect ratio is reached, the critical motile pressure to unjam is predicted to scale as 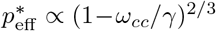 (supplementary eq. S4). Here, the sub-linear 2*/*3-scaling with the relative cell-cell tension 1 − *ω*_*cc*_*/γ* emerges from adhesive tension also increasing the vertical aspect ratio of the cell (see Fig. 1b), which reduces the elongation due to the dipole. Since this analysis neglects the contribution of rotational diffusion, it is only valid when the persistence time is sufficiently large, which is the case in our simulations, where *τ*_*p*_ *≫η*_*c*_*t*_*c*_*/γ*, supplementary Fig. S2.

The predicted scaling aligns well with observed neighbor exchange rates in simulations, Fig. 3d. As expected, cells are more elongated when they are unjammed in confluent tissue, Fig. 3e. When measuring 2D cell shape, the critical shape index 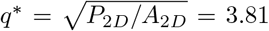, originally obtained from vertex models, provides an excellent delineation of the solid-to-fluid transition in the confluent regime, Fig. 3f. In our model, active cell elongation is driven by two potentially independent factors. First, the force dipole *p*_*a*_*/*(*λ*_*a*_*L*_*c*_) induces elongation along the cell’s polarity while remaining static, analogous to the preferred perimeter in self-propelled Voronoi models [20]. Second, the net migratory pressure *p*_*a*_ drives net migration, similar to the self-propulsion velocity in Voronoi models, while also elongating cells due to substrate friction and resistance from neighboring cells. Despite their distinct effects on single-cell dynamics, both parameters regulate the solid-to-fluid transition through the same critical shape index, supplementary Fig. S3 and supplementary Fig. S4.

### D. Pull-off force controls tissue fluidity at low adhesive tension

At low adhesive tension, we observe that adhesive tension promotes tissue jamming, a behavior that was previously linked to the existence of intercellular pores [23] (Fig. 3b,d) At high *p*_eff_, intercellular pores emerge as cells delaminate in response to local crowding, effectively reducing the density of the bottom layer. However, we also observe fluid-like behavior in fully confluent tissue (see supplementary Fig. S5). We propose that this behavior arises from the inclusion of normal adhesive forces and cell-cell debonding dynamics in our model. Here, adhesion energy acts as an energy barrier for T1 transitions. Hence, we expect the critical activity to unjam to scale as 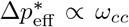 (supplementary eq. S5), where 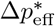 denotes the shift in critical activity relative to the non-adhesive (density dependent) jamming point. Notably, because this regime is dominated by debonding rather than cell elongation, it can give rise to fluid-like behavior even below the critical shape index, similar to subconfluent states (Fig. 3f).

At high number density, the density-dependent jamming transition is a dominant mechanism at low adhesive tension. To better understand the role of adhesive tension in driving solidification in this regime, we simulated porous tissues at lower number density 0.6. Here, increasing adhesive tension *ω*_*cc*_*/γ* drives cell movements from a diffusive to a subdiffusive regime (Fig. 4a). Similarly, the diffusivity *D*_eff_ decreases by orders of magnitude by increasing cell-cell adhesive tension and by decreasing motile pressure Fig. 4b, indicating caging by neighboring cells. Caging occurs when motility-driven agitation is too weak to overcome energy barriers to neighbor exchanges, which depend on cell density and adhesive tension (Fig. 4c). Therefore, we simulated the minimal motile pressure *p*_eff,*S*_ that is needed for cell separation, Fig. 4d, which follows a similar trend to the BWdG relation, with the separation force increasing linearly with adhesive tension *ω*_*cc*_ at low *ω*_*cc*_. We find that this solid-to-fluid transition, measured by the rate of neighbor exchanges, follows the scaling of the pull-off force (Fig. 4e). This is consistent with a model in which tissue fluidity is governed by cell-cell separation, with adhesion promoting a jammed or glassy state, similar to attractive colloidal systems [51].

**FIG. 4.**
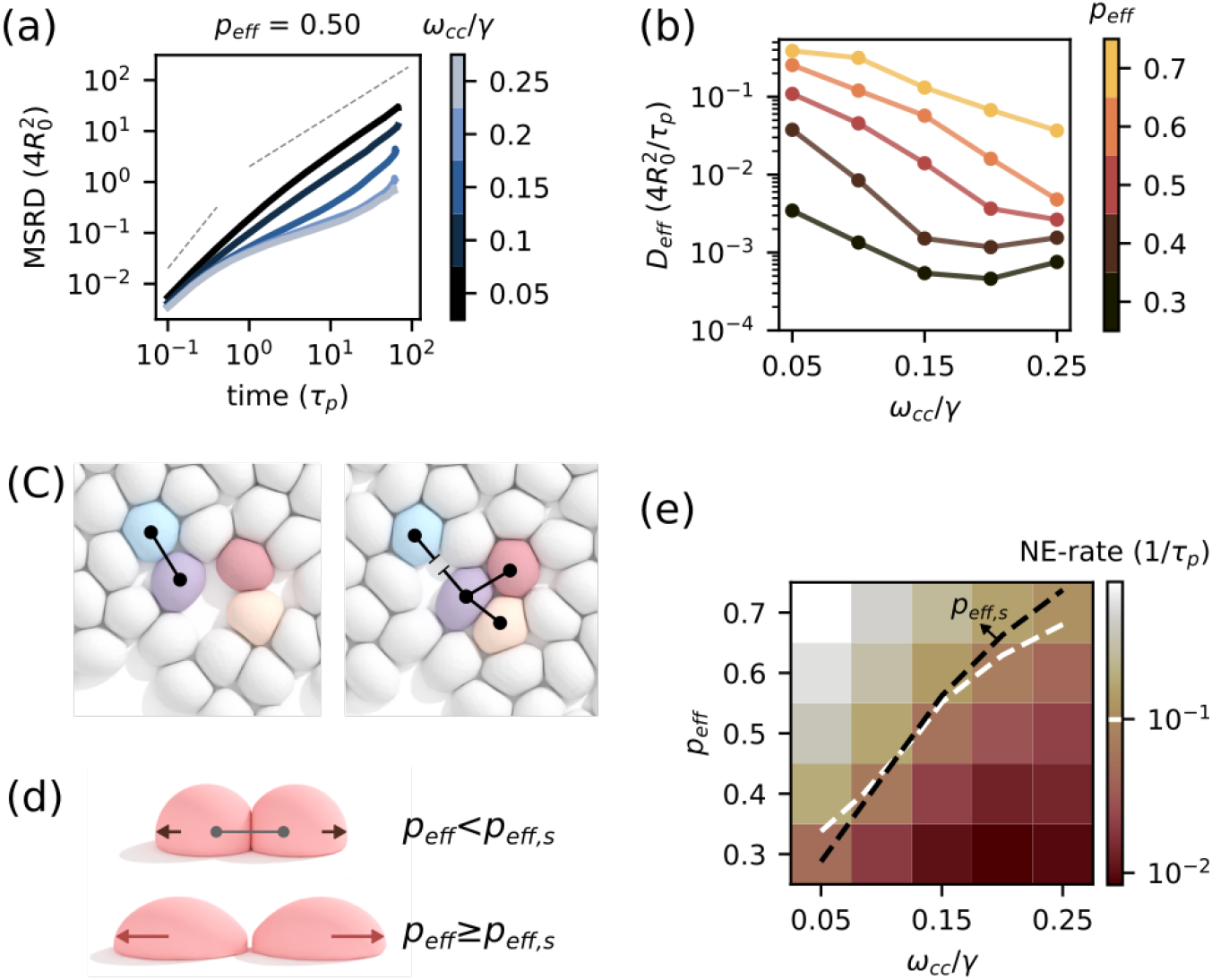
Fluidity of porous tissue at number density 0.6. (a) Mean squared relative displacement (MSRD) of simulated tissue in function of cell-cell adhesive tension *ω*_*cc*_. Dotted lines indicate super diffusive and diffusive limits. (b) Normalized effective diffusion constant *D*_eff_ = lim_*t*→∞_ MSRD(*t*)*/*4*t* for changes in activity *p*_eff_ and adhesive tension *ω*_*cc*_. (c) Example of cell-cell separation in porous tissue. (d) Minimal activity *p*_eff,7_ needed for separation of two cells migrating away from the cell-cell contact. (e) Measurement of neighbor exchange rate. We set a threshold NE-rate = 10^−1^(1*/τ*_*p*_) distinguishing fluid-like from solid-like tissue indicated by the white dashed line. The black dashed line indicates *p*_eff,*S*_. Note that this pull-off force is not linear with adhesive tension *ω*_*cc*_. This is expected since in our cell motility model we assume cell migration enhances cell elongation (and decreases *R*_*a*_), therefore decreasing the pull-off force as predicted by BWdG. See also supplementary Fig. S6 and supplementary video S2 for a visualization of the simulated tissues.

### E. Cell-cell friction as a distinct mechanism from jamming in monolayers

Finally, we investigate the effect of cell-cell friction *ξ*_*cc*_ on tissue dynamical properties. We simulate an active confluent tissue at *ω*_*cc*_ = 0.5, number density of 1.1, and vary *ξ*_*cc*_*/ξ*_*cs*_, the ratio of cell-cell to cell-substrate friction. Increasing intercellular friction drives tissue dynamics from diffusive to sub-diffusive behavior (Fig. 5a) and greatly decreases neighbor exchange rates (Fig. 5b), effectively appearing to jam the tissue. This jamming can be explained by the slow-down of T1-transition dynamics due to friction, where the tissue will jam when the persistence time *τ*_*p*_ becomes smaller than the timescale of T1-transitions. This observation is in line with recent results from phase field model simulations [35] and experiments which observed jamming associated with maturing cell-cell adhesion proteins [52]. Remarkably, however, the critical cell shape index *q*^∗^ = 3.81 does not align with the transition predicted by the NE-rate (Fig. 5c), hinting at a distinct mechanism from shape-controlled jamming in the vertex model. To clarify this discrepancy, we examine the rheological properties of the tissue through a simulated shear experiment, where an instantaneous shear strain *ϵ*_*xy*_ is applied to an active tissue, and the resulting shear stress *σ*_*xy*_ is tracked over time (Fig. 5d). We observe a relaxation consisting of two phases: a first fast relaxation, where T1 transitions occur through external shear stress on the cells, and a second slow relaxation where cell-scale migration forces may overcome remaining local energy barriers of T1-transitions. Notably, due to the applied shear stress, the motile pressure *p*_eff_ required to fluidize the system is expected to be lower than in the absence of shear [53, 54]. This relaxation can be approximated by a stretched exponential [23] with a relaxation timescale *τ*_*R*_ and a residual stress *σ*_*Y*_, i.e., the stress that remains in the tissue after relaxation. This residual stress can be interpreted as the minimal stress for the tissue to plastically deform: the yield stress. As expected from our previous results (Fig. 3 and supplementary Fig. S8), this yield stress vanishes by increasing cell activity (Fig. 5e,g), thus driving the tissue to a true fluidized state. On the other hand, the relaxation timescale *τ*_*R*_ does not change with cell activity, since this timescale primarily represents short passive relaxation. In contrast, we find that intercellular friction does not affect the yield stress, but only the relaxation timescale *τ*_*R*_ (Fig. 5f-h) suggesting that only the viscous properties of the tissue change with cell-cell friction. Thus, tissues may appear solid-like based on cell dynamics alone, even though friction does not affect their static load-bearing capacity. Cell dynamics alone are therefore insufficient to distinguish true solid-from liquid-like states.

**FIG. 5.**
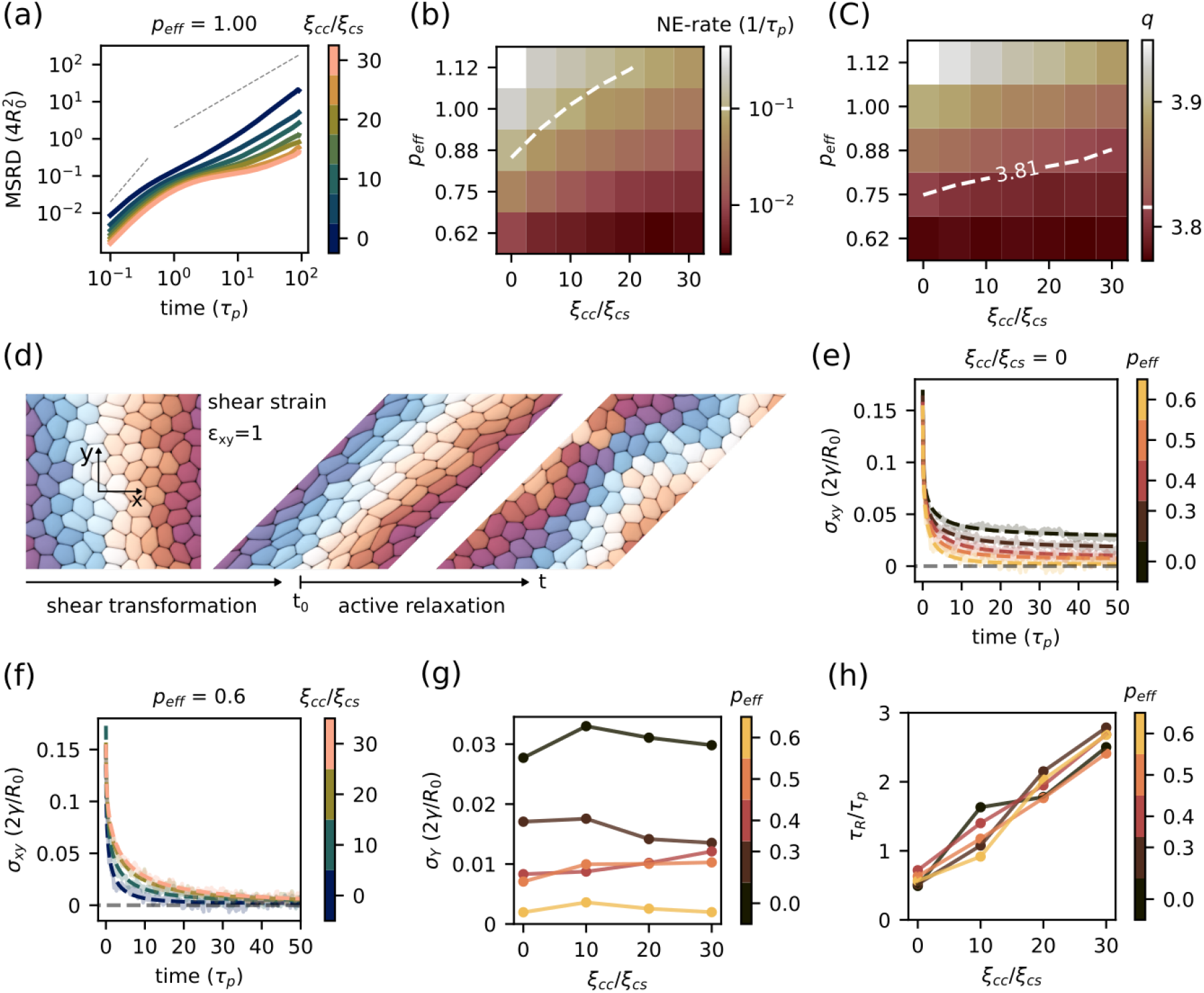
The effect of cell-cell friction on tissue fluidity. (a) MSRD in function of friction ratio *ξ*_*cc*_*/ξ*_*cs*_. (b) Neighbor exchange rate separating jammed from fluid-like tissue at the threshold NE-rate = 10^−1^(1*/τ*_*p*_). (c) Cell shape index *q* where the line indicates the critical shape index predicted by vertex models *q*^#x002A;^ = 3.81. See supplementary Fig. S7 and supplementary video S3 for visualization of the simulated tissues. (d) Example of a tissue deformed by a shear strain *ϵ*_*xy*_ = 1. After this instantaneous deformation, the tissue relaxes by passive and active T1-transitions. Individual cells are colored in distinct colors as a visual guide. (e-f) Stretched exponential fit of shear stress *σ*_*xy*_ during relaxation in function of (e) activity *p*_eff_ and (f) friction ratio. (g) Yield stress *σ*_*Y*_ and (h) relaxation time *τ*_*R*_ in function of friction ratio and cell activity.

## III DISCUSSION

In this work, we presented a novel model of active cell monolayers based on deformable cells with explicit contact interactions, parameterized by physical properties of the acto-myosin cortex. With this model, we examined the distinct mechanisms by which adhesive tension regulates cell dynamics and governs tissue mechanical properties [55].

On the one hand, adhesive bond strength governs the pull-off force between neighboring cells, reducing rearrangement rates and thereby decreasing tissue fluidity. This gives rise to a regime in which the critical activity required to fluidize the tissue increases linearly with adhesive tension (Fig. 6). Such behavior dominates in porous tissues, consistent with observations of mesodermal fluidization during embryogenesis [5, 6, 17]. However, this regime also may emerge in confluent tissues — despite the absence of intercellular pores — when activity is sufficient to overcome density-dependent jamming. In this case, tissues may appear fluid-like even with a shape index below the critical value predicted by vertex models. On the other hand, adhesion proteins like E-cadherins are linked to the cytoskeleton, thereby downregulating actomyosin tension and effectively reducing interfacial tension [30]. This expands cell-cell contacts and promotes cell elongation through the reduction of the net tension at the cell-cell interface. Our results demonstrate that, in confluent tissue, decreased interfacial tension enhances tissue fluidity by facilitating cell deformation and elongation, in line with observations of primary human bronchial epithelial cells [13] and vertex model predictions [19, 20] (Fig. 6). However, the mechanism underlying cell elongation is different compared to these models. In vertex models, the preferred cell shape *q*_0_ itself is an input model parameter that depends on interfacial tension *γ*_*c*_ and actomyosin contractility with elasticity *κ* (as *q*_0_ = − *γ*_*c*_*/*2*κ*) [19]. In this approach, cell elongation can be achieved by negative interfacial tension. In contrast, we modeled cells with positive tension, consistent with force inference studies of epithelial tissue [56] and current understanding of interfacial actomyosin tension [30]. Rather, elongated shapes result from extracellular forces (cells pushing neighboring cells) and active intracellular (static and dynamic) force dipoles. For these active mechanical properties, we were able to derive and verify explicit scaling laws for the fluid-to-solid transition, both for low and high adhesive tension. The interplay of these two regimes gives rise to a phase diagram that bridges unjamming in attractive colloidal systems and shape-dependent unjamming at high adhesive tension, as shown in Fig. 6.

**FIG. 6.**
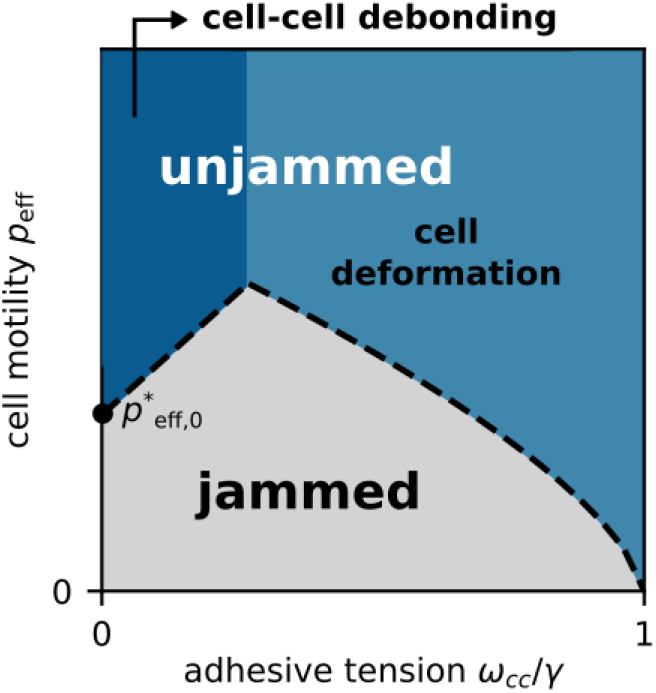
Schematic representation of the jamming phase diagram in function of adhesive tension and cell motility based on Fig. 3d. We consider two distinct mechanisms of tissue fluidization: cell-cell debonding (colloidal limit) or cell deformation (vertex limit). In the low adhesive regime fluidization is limited by cell-cell debonding, where adhesion induces solidification. In the high adhesive regime fluidization is limited by cell deformation, where adhesive tension promotes fluidiza-tion. Here, 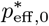 indicates the density dependent jamming transition for non-adhesive particles. Note that at point *J*, the density to unjam at zero adhesive tension and activity, 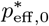 will move towards the origin.

Finally, with our model, we were able to investigate the role of cell-cell friction, which may emerge from adhesion turnover dynamics at long timescales [33, 34]. These dynamics have been proposed to affect epithelial tissue mechanical properties during embryogenesis [55, 57] and in-vitro cell culture [52]. Our results show that intercellular friction slows tissue dynamics, creating an apparent jamming effect despite a vanishing yield stress, indicating the absence of true solid-like behavior. However, cell shape alone does not differentiate between diffusive and arrested dynamics due to cell-cell friction: At high cell-cell friction, cells can deform beyond the critical shape while still displaying subdiffusive movements. Notably, this is mirrored in various observations in epithelial tissue, where the shape index remains above the critical value, yet cell dynamics exhibit jammed behavior [13, 58–60]. In other cases, however, vertex model predictions hold [61]. These discrepancies may stem, at least in part, from the omission of intercellular friction in vertex theory.

In addition, these results can help in understanding the role of (partial) epithelial-to-mesenchymal transition (pEMT) that has been associated with increased invasiveness and metastasis during tumor progression. Although it is not yet clear whether the unjamming transition (UJT) and pEMT are similar [62, 63], both pEMT and UJT involve an initially solid-like epithelial tissue that fluidizes and remodels. The UJT in epithelial tissue is often characterized by sustained E-cadherin expressions and increased adhesiveness, related to shape changes, maintaining epithelial barrier function. In contrast, during pEMT, cells lose E-cadherin-mediated adhesions, cell polarity and increase their migratory potential, whereas vertex models would predict solidification by decrease of adhesiveness. This model offers two possible explanations: In the low adhesive regime, decrease of adhesion will fluidize the tissue, similar to colloidal systems, even at confluent density. An alternative explanation could lie in differences in bond kinetics, since N-cadherins and P-cadherins, associated with mesenchymal cell types, show faster turnover compared to E-cadherin [30, 64]. This would effectively decrease intercellular friction and result in increased cell dynamics. However, the exact effects of E-cadherin on cell shape are still not well known. For example, both upregulation and downregulation of E-cadherin can lead to cell elongation and tissue fluidization [55, 65]. Hence, the precise effects of intracelular mechanobiological factors such as adhesive ligands on cell mechanical properties - and consequently on tissue rheology - remain to be further elucidated.

This work highlights the importance of cell-cell contact interactions in understanding the (un)jamming transition in biological tissue. By explicitly accounting for intercellular tension, cell-cell debonding dynamics, and cell-cell friction, we bridge the gap between colloidal-like and vertex-based models of tissue mechanics. Moving forward, incorporating biased polarization mechanisms — such as cell-cell alignment or contact inhibition of locomotion — could shed light on their role in tissue dynamics and collective migration during embryogenesis and cancer metastasis [66]. This modeling framework also offers a platform to study how mechanical cell heterogeneity affects tissue rheology, providing insights into tumor mechanics, where cell heterogeneity is one of the hallmarks of cancer [67]. Furthermore, cell populations during embryonic development are often heterogeneous, involving phenomena such as cell mixing and cell sorting. In these contexts, the role of tissue (un)jamming still remains an open question.

## IV METHODS

### A. Active foam model

Tissue structure is determined by the balance between interfacial and cell-medium surface tension, similar to a liquid foam, but also active forces from acto-myosin tension fluctuations, cell divisions and cell migration [23, 45, 68]. We model a cell monolayer as an active foam, representing individual cells as pressurized bubbles with a resting radius *R*_0_ when detached, Fig. 1. We assume that cell shape is governed by the actomyosin cortex, which we assume to behave as a curved 2D liquid film at long timescales, with constant viscosity *η*_*c*_. For simplicity, we assume equal shear viscosity, longitudinal, and bulk viscosity of the cortex. Hence, the total in-plane cortex tension is

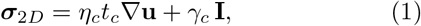

with in-plane velocity **u** = (*u, v*), and where surface tension *γ*_*c*_ = *γ* at free surfaces, and *γ*_*c*_ = *γ* − *ω*_*cj*_ at contacts with surface *j*. This accounts for the reduction of actomyosin contractility at cell-cell (and cell-substrate) interfaces [30], producing an effective adhesive tension *ω*_*cj*_. The in-plane tension balance then reads

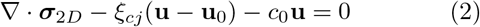

where the second term includes cell-cell and cell-substrate friction with friction coefficient *ξ*_*cj*_ [69], and cell-medium friction with damping constant *c*_0_. The relative contribution of friction compared to cortex viscosity is reflected in the hydrodynamic length-scale *L*_*η,j*_ = (*η*_*c*_*t*_*c*_*/ξ*_*cj*_)^1*/*2^, with typically *L*_*η,s*_ *< R* [70]. It should be noted that *L*_*η,s*_ is distinct from the tissue-scale hydrodynamic length *L*_*h*_ = (*η*_*t*_*/ξ*_*cs*_)^1*/*2^, which depends on the (emergent) 2D tissue viscosity *η*_*t*_ and typically spans multiple cell diameters [71]. Since this stress acts on a curved surface, out-of-plane forces arise. Assuming a Kirchoff-Love approximation of viscous bending, and with *w* the out-of-plane velocity, we write

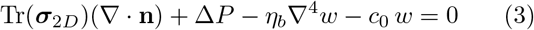

with viscous bending constant 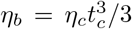, and where Δ*P* = *P*_*b*_ + *P*_*c*_ + *P*_*a*_ is the pressure drop over the surface that is caused by the internal pressure *P*_*b*_, contact pressure *P*_*c*_, and an active protrusive pressure *P*_*a*_. Note that this equation reduces to the Young-Laplace law for the equilibrium shape (*w* = 0). We assume the cell cytoplasm behaves as a near-incompressible fluid. Hence, the internal pressure *P*_*b*_ serves as a Lagrange multiplier to ensure incompressibility. The contact pressure *P*_*c*_ includes an adhesive and a repulsive term,

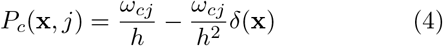

where *δ*(**x**) is the overlap distance between contacting surfaces at position **x** and where *h* the effective range of the adhesive contact with surface *j*. Cell migration is modeled by a force dipole, causing cell elongation, and a net protrusive force in the direction of the cell polarization **p**, causing cell displacements. The active pressure on the cell surface at position **x** is

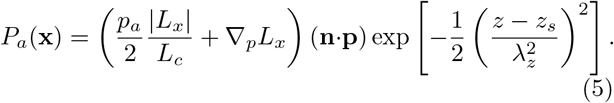

Here, *L*_*x*_ = (**x − x**_0_) **p** is the distance from the surface to the cell center **x**_0_ in the direction of polarization. Furthermore, we assumed both net migration and cell elongation scale by the same active parameter *p*_*a*_ such that _*p*_ = *p*_*a*_*/*(*λ*_*a*_*L*_*c*_), where *L*_*c*_ is the instantaneous radius of the cell in the direction of polarization and *λ*_*a*_ is the dipole scaling factor: smaller *λ*_*a*_ implies a stronger dipole. Furthermore, we assume that the cell migrates through acto-myosin activity close to the cell’s substrate. Therefore, we localize the protrusive forces in a narrow region near the substrate, located parallel to the *xy* plane at position *z* = *z*_*s*_), by scaling *P*_*a*_(**x**) with a Gaussian kernel with length scale *λ*_*z*_ similar to [72]. The cell activity *p*_*a*_ is adimensionalized by balancing the active pressure with the Young-Laplace pressure from free surface tension *γ* (see SI) as,

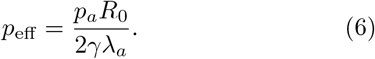

The cell polarity vector **p** = (cos *ϕ*, sin *ϕ*, 0) is a minimal representation of the cell front-to-rear polarity, restricted to the substrate plane *xy*. The cell polarity undergoes rotational diffusion

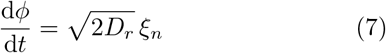

where *ξ*_*n*_ is standard, Gaussian noise and *D*_*r*_ the rotational diffusion coefficient corresponding to persistence time *τ*_*p*_ = 1*/D*_*r*_.

To simulate the dynamics of multicellular systems, we use a particle-based model where the cell surface is discretized in a triangulated mesh, Fig. 1a. At each time step, mechanical forces acting on the vertices of the mesh are computed, and vertex displacements are obtained by integrating the overdamped equation of motion in the semi-implicit system **Λ u** = **F**, where **Λ** is a grand resistance matrix containing the friction and viscous elements, and **F** all other forces. Details of the numerical discretization and computational implementation of this model are provided in the Supplementary Material (see also references [30, 45, 47, 72–76] therein).

### B. Mean squared relative displacement

To exclude the effects of global tissue movements, we compute the relative 2D distances **x**_*ij*_ = **x**_*i*_ − **x**_*j*_ between cell pairs *i, j*. The mean squared displacement of these relative distance is then computed for lag times Δ*t* as

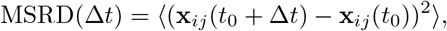

where the cell pairs are in contact at time *t*_0_, and ⟨⟩ denotes the average over both cell pairs and time. The MSRD is normalized by the cell diameter MSRD(Δ*t*)*/*(2*R*_0_)^2^.

### C. Neighbor exchange rate

As a measure of T1-transitions, the NE-rate is computed as the total number of new cell-cell contacts observed over the persistence time *τ*_*p*_ averaged over all cells visualized in Fig. 3d and Fig. 4d.

### D. Virial stress

The stress on tissue level is estimated by the virial stress theorem [77, 78] as

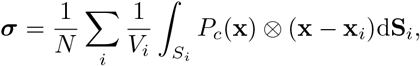

where *N* is the number of cells, *P*_*c*_(**x**) is the contact pressure from neighboring cells on cell *i*, **x** is the contact point, **x**_*i*_ is the cell center, **S**_*i*_ is the directed cell surface pointing outwards, and *V*_*i*_ is the cell volume. This stress is smoothed over time by a moving average with a window size of *τ*_*p*_*/*10, to limit the effect of fluctuating contact forces due to the discrete nature of the contacts. The hydrostatic tissue stress is then given by *σ*_*h*_ = tr(***σ***)*/*3.

### E. Cell shape index

The apical cell shape is computed by projecting the cell nodes on the substrate. The convex hull of these projected points is computed. From the convex hull, the perimeter *P*_*i*_ and projected area *A*_*i*_ are calculated for each cell i. The shape index for each cell is then given by 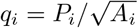.

### F. Tissue density

We measure 2D tissue density by projecting the cells on the substrate. The total area of this projected tissue is divided by the domain size *A*_box_. This density differs from the number density, which is defined as 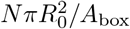, where *N* is the total number of cells with cell volume 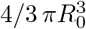.

### G. Simulation setup

*N*_0_ cells with radius *R*_0_ are seeded in a periodic boundary domain in a hexagonal pattern, where cell positions are displaced by a distance from a Gaussian distribution with standard deviation *R*_0_. Next, a fraction of 0.35*N*_0_ cells are removed using a Latin hypercube sampling method to ensure a uniform distribution of *N* cells in the periodic domain. In all simulations, the tissue exists of *N* = 100 cells, unless otherwise specified. Next, the size of the domain decreases linearly with time with velocity *v*_*b*_ until the desired number density 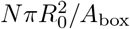 is reached, with *A*_box_ the final size of the periodic domain, after a time *τ*_init_ = 360*η*_*c*_*t*_*c*_*/γ*. Simultaneously, a centripetal force 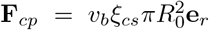 is added to all cells during *τ*_init_. Cells are allowed to relax for an additional time of *τ*_init_*/*3, without any active forces. These monolayers are then used to initialize the tissue simulations. For simulations shown in Fig. 2, the cell active migration pressure exponentially relaxes in time as *p*_eff_ (*t*) = exp(− 2*t/*(3*τ*_*p*_)). This effectively allows the system to cool down to an annealed state. For all other simulations, the cell activity *p*_eff_ is set fixed, and the tissue is allowed to reach steady-state. Measurements of tissue dynamics only started after an additional simulation time of 2*τ*_*p*_. The model parameters that were used for the simulation can be found in supplementary Table S1. All simulations were conducted in Python using the Mpacts particle-based simulation framework.

## DATA AVAILABILITY

The simulation scripts and framework are available on gitlab: https://gitlab.kuleuven.be/mebios-particulate/publications/rigidity_transitions_cell_monolayers.git [79].

## ACKNOWLEDGMENTS

J.V. acknowledges support from the Research Foundation Flanders (FWO), grant 11D9923N. B.S. acknowledges support from the Research Foundation Flanders (FWO), grant 12Z6118N, and KU Leuven internal funding C14/18/055.

J.V., H.R. and B.S. conceptualized the work; J.V. performed the simulations; J.V. performed the analysis; J.V. and B.S. wrote the manuscript; and all authors edited the manuscript.

## SUPPLEMENTARY TABLES

**Supplementary Table S1.**
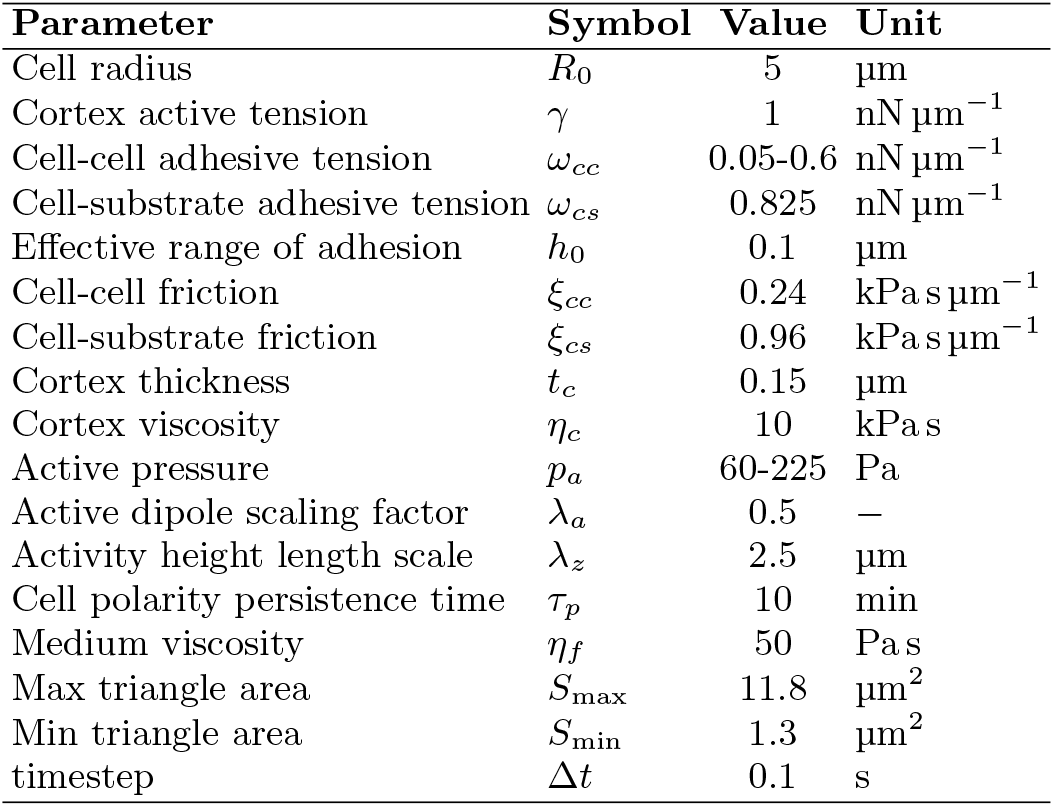
model parameters

## SUPPLEMENTARY VIDEOS

Supplementary Video S1. Dynamic tissue at number density 1.1, *p*_eff_ = 0.875, and varying adhesive tension *ω*_*cc*_*/γ* for a simulation time 50*τ*_*p*_.

Supplementary Video S2. Dynamic tissue at number density 0.6, *p*_eff_ = 0.5, and varying adhesive tension *ω*_*cc*_*/γ* for a simulation time 50*τ*_*p*_.

Supplementary Video S3. Dynamic tissue at number density 1.1, *ω*_*cc*_*/γ* = 0.5, *p*_eff_ = 0.875, and varying cell-cell friction *ξ*_*cc*_*/ξ*_*cs*_ for a simulation time 50*τ*_*p*_.

## SUPPLEMENTARY FIGURES

**Supplementary Fig. S1.**
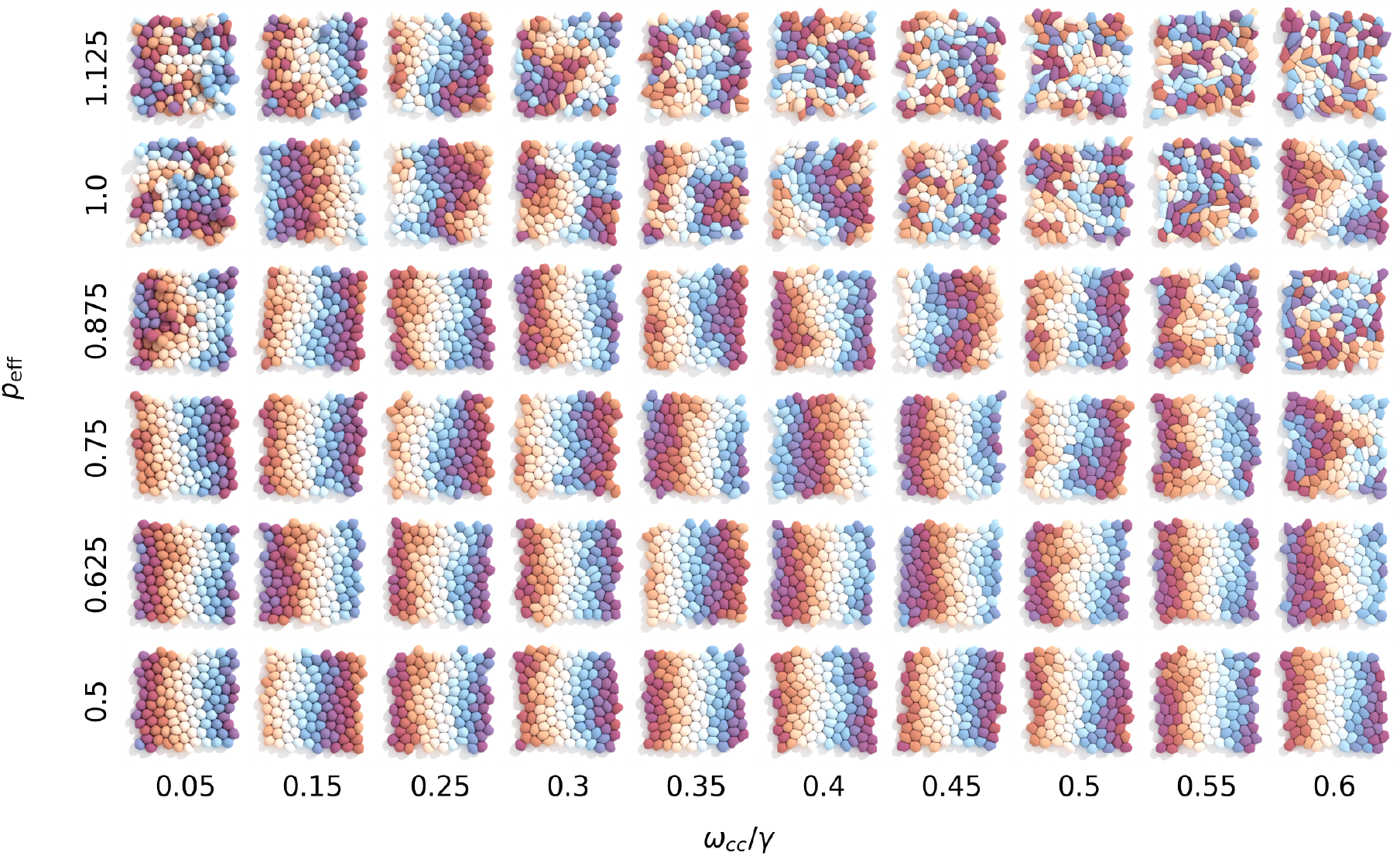
Snapshot of tissue at *t* = 50*τ*_*p*_ for confluent tissue (number density 1.1), varying cell motility *p*_eff_ and adhesive tension *ω*_*cc*_*/γ*, where color indicates the initial position of the cell. Notice that cells at low *ω*_*cc*_*/γ* and high *p*_eff_ often delaminate from the tissue that effectively reduces cell density.

**Supplementary Fig. S2.**
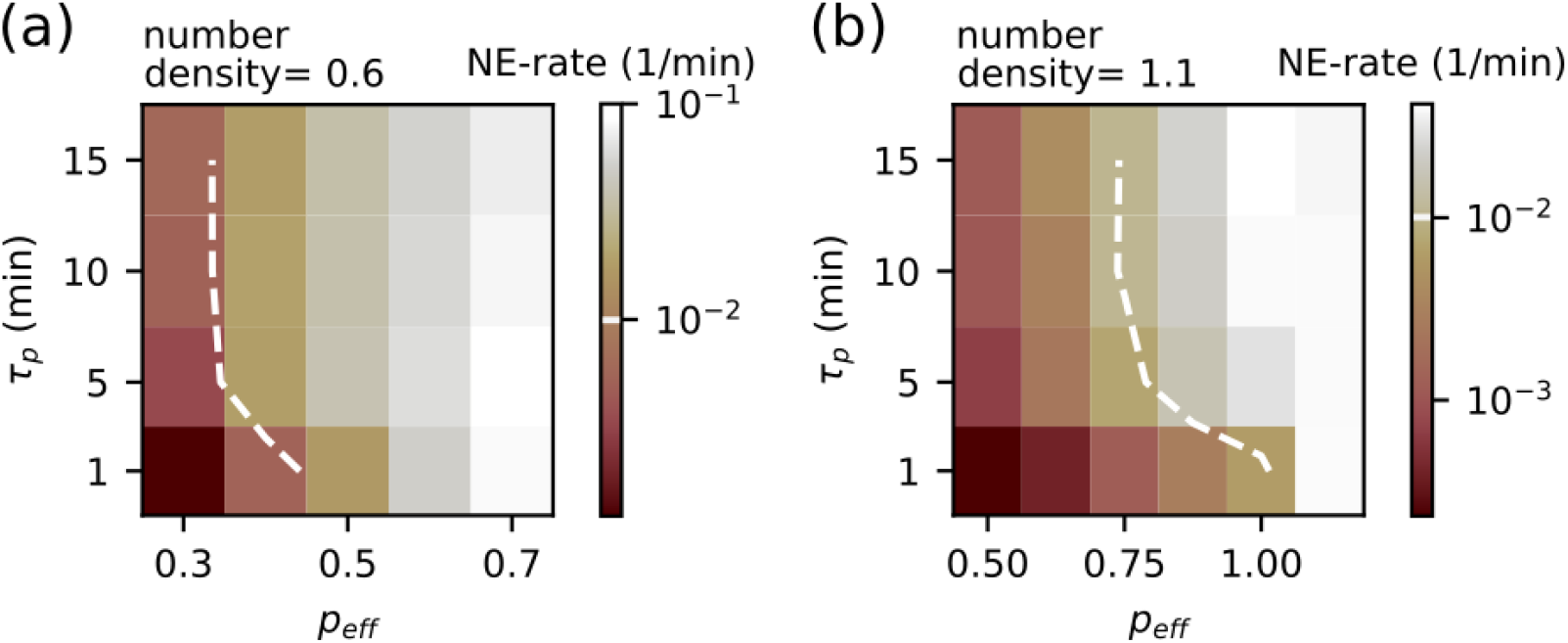
Effect of persistence time *τ*_*p*_ on neighbor exchange rate (NE-rate) (cell dynamics) in (a) porous tissue for number density = 0.6, and adhesive tension *ω*_*cc*_*/γ* = 0.05 and (b) confluent tissue for number density = 1.1, and adhesive tension *ω*_*cc*_*/γ* = 0.5. The NE-rate increases with *τ*_*p*_ and saturates after a persistence time of *τ*_*p*_ ≥ 5(min). We set the solid-fluid threshold at NE-rate = 10^−2^ (1*/*min).

**Supplementary Fig. S3.**
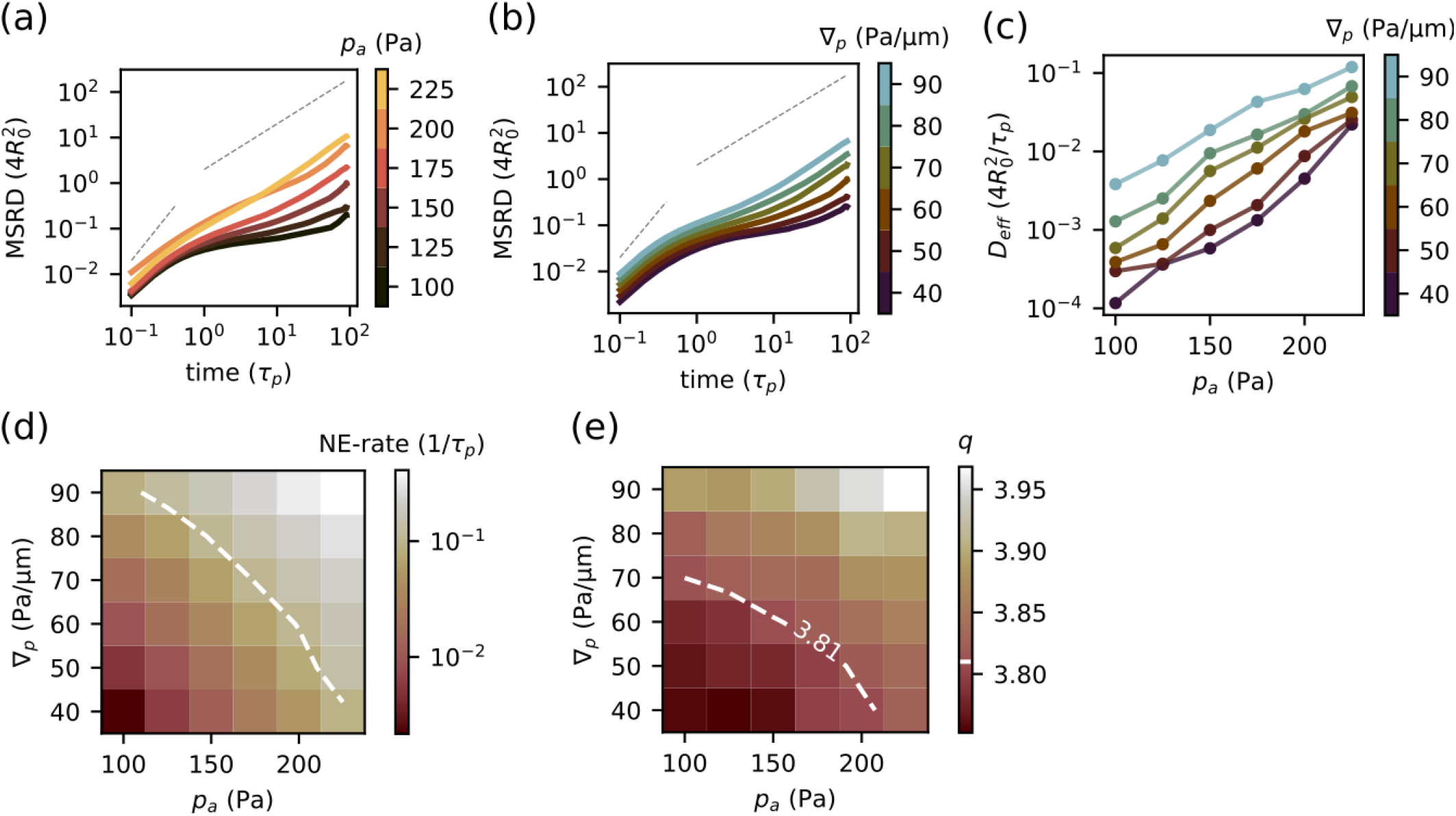
(a) MSRD in function of migration pressure *p*_*a*_, (b) and in function of migration dipole ∇_*p*_. Both *p*_*a*_ and ∇_*p*_ can shift the tissue dynamics from subdiffusive (indicative of caging) to diffusive behavior. (c) Effective diffusion coefficient normalized by cell radius *R*_0_ and persistence time *τ*_*p*_. (d) Similarly, the neighbor-exchange rate changes orders of magnitude in function of *p*_*a*_ and ∇_*p*_. We set a threshold at NE-rate = 10^−1^(1*/τ*_*p*_) above which we quantify tissue as fluid-like. (e) We show that the critical cell shape index *q*^∗^ = 3.81, indicated by the dashed line, follows the same trend as the NE-rate. In other words, independent on the means of cell deformation – by internal or external forces – tissue fluidity indeed can be predicted by cell shape as expected from vertex theory.

**Supplementary Fig. S4.**
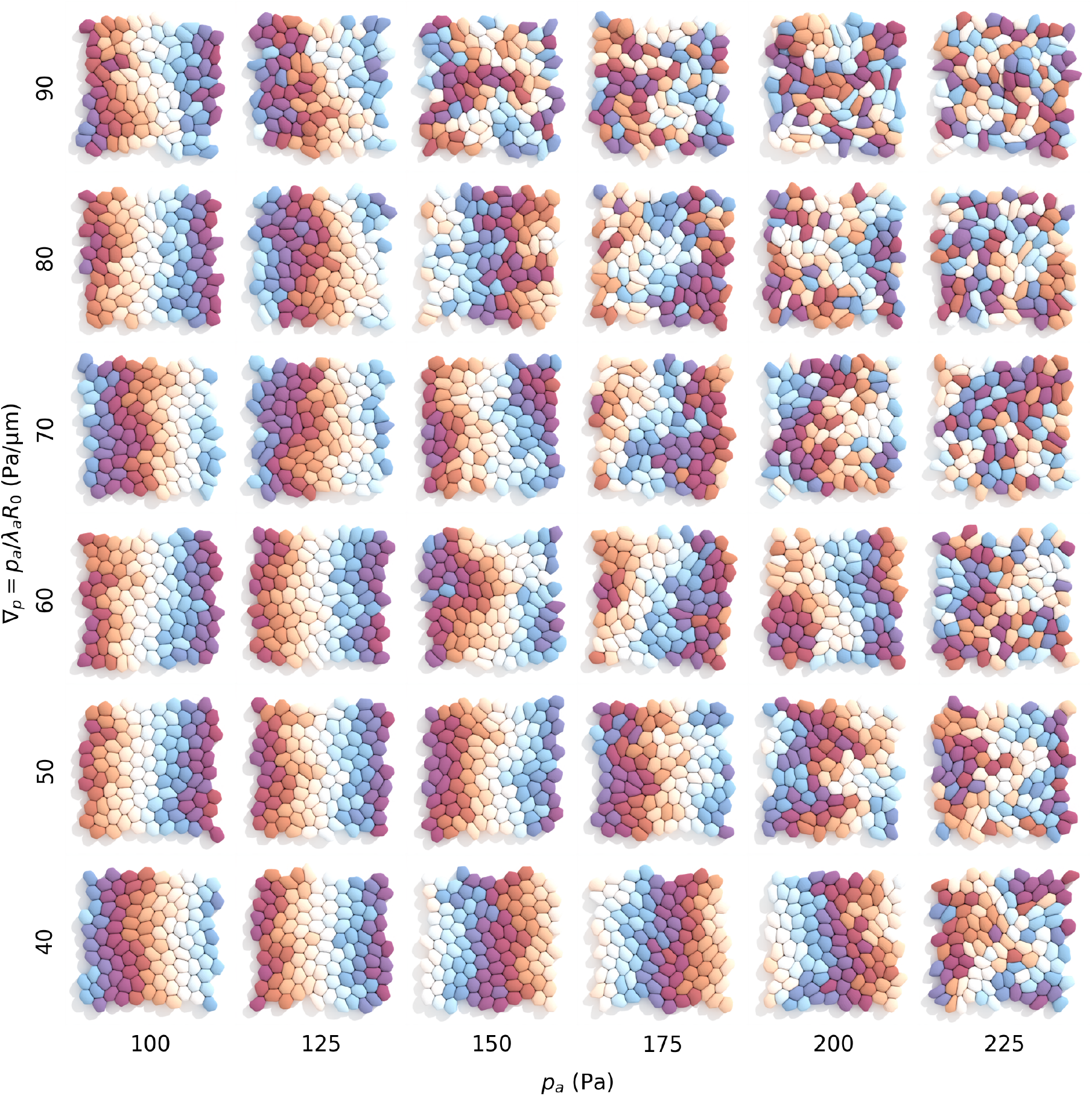
Snapshot of tissue at *t* = 50*τ*_*p*_ for confluent tissue (number density 1.1, *ω*_*cc*_*/γ* = 0.5), varying cell dipole ∇_*p*_ and migration pressure *p*_*a*_, where color indicates the initial position of the cell.

**Supplementary Fig. S5.**
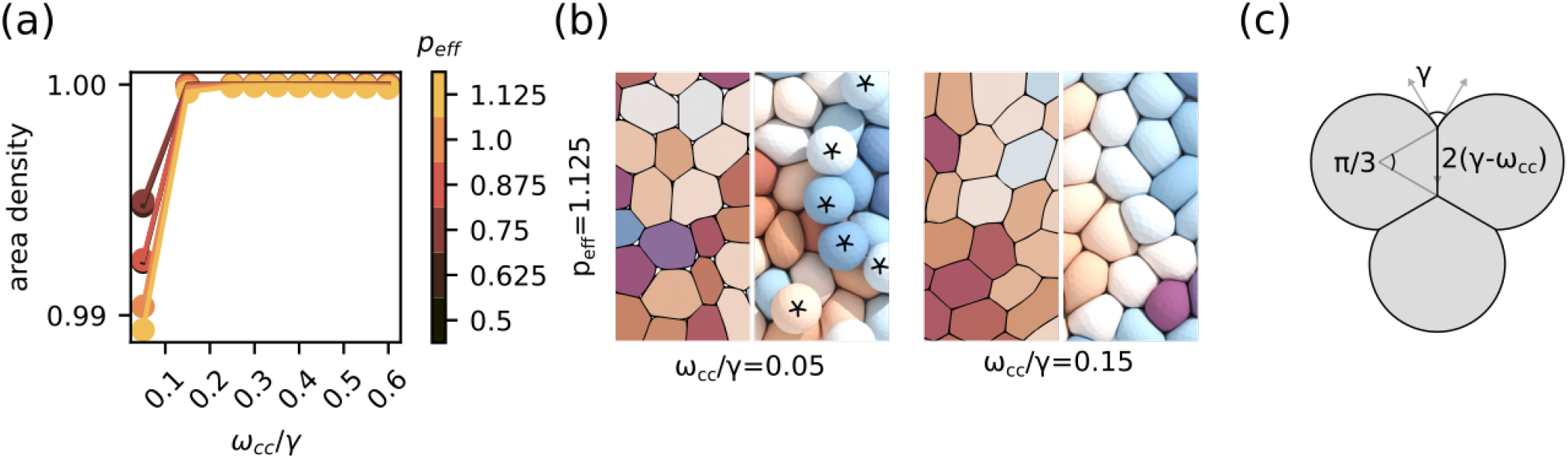
(a) Observed area density of tissue at high number density (= 1.1) varying adhesive tension *ω*_*cc*_*/γ* and activity *p*_eff_. (b) Snapshot of tissue at low *ω*_*cc*_*/γ* and high *p*_eff_, where each individual cell is assigned a distinct color for visual clarity. Cells delaminate at *ω*_*cc*_*/γ* = 0.05 labeled by an asterisks and tissue becomes microporous. (c) Assuming an adhering cell triplet, the minimal adhesive tension needed to close the pore is given by *ω*_*cc*_*/γ* ≥ 1 − cos (*π/*6). Therefore, it is expected that microporosity arises for *ω*_*cc*_*/γ <* 0.134.

**Supplementary Fig. S6.**
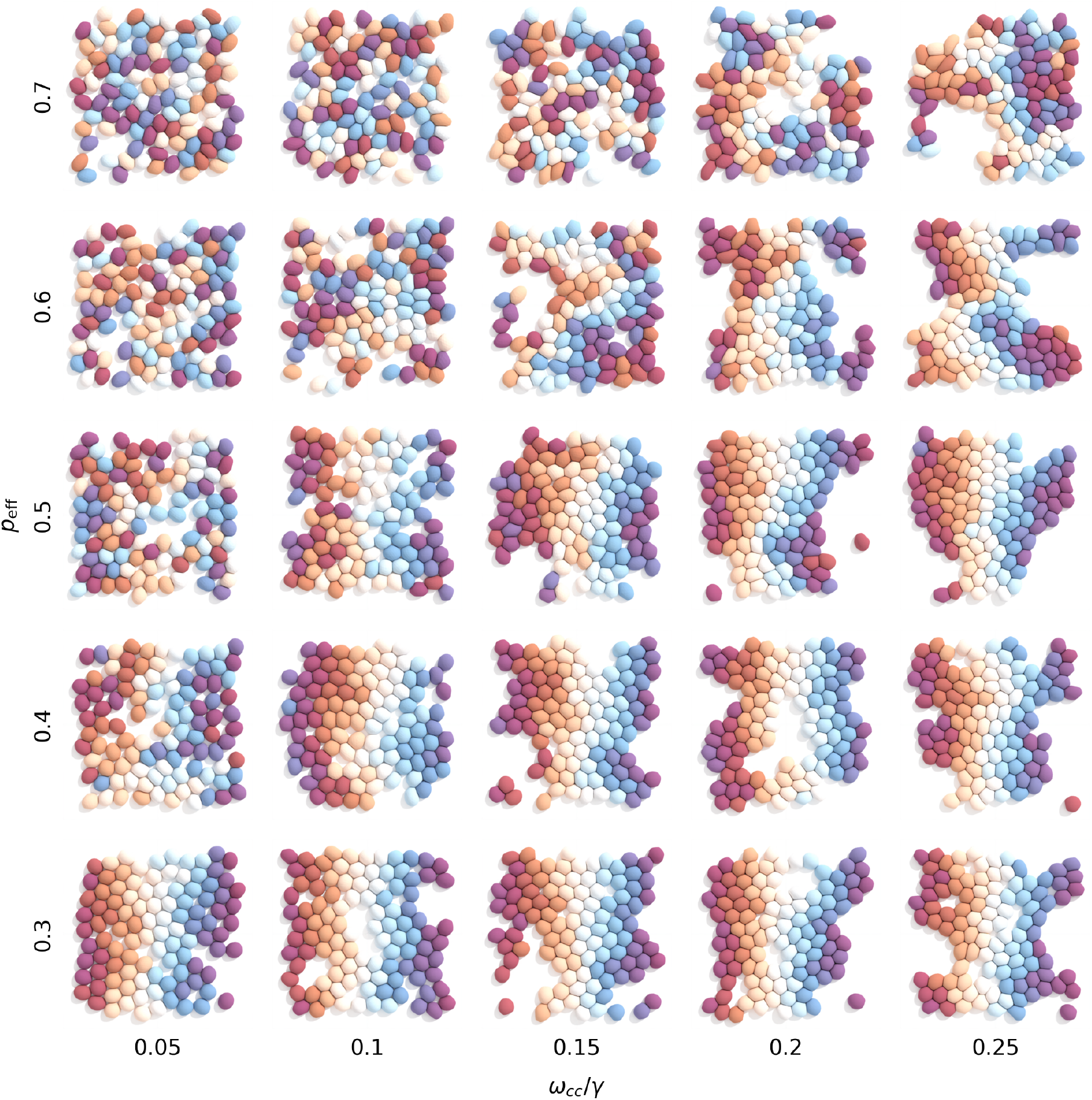
Snapshot of tissue at *t* = 50*τ*_*p*_ for porous tissue (number density 0.6), varying cell motility *p*_eff_ and adhesive tension *ω*_*cc*_*/γ*, where color indicates initial position of the cell.

**Supplementary Fig. S7.**
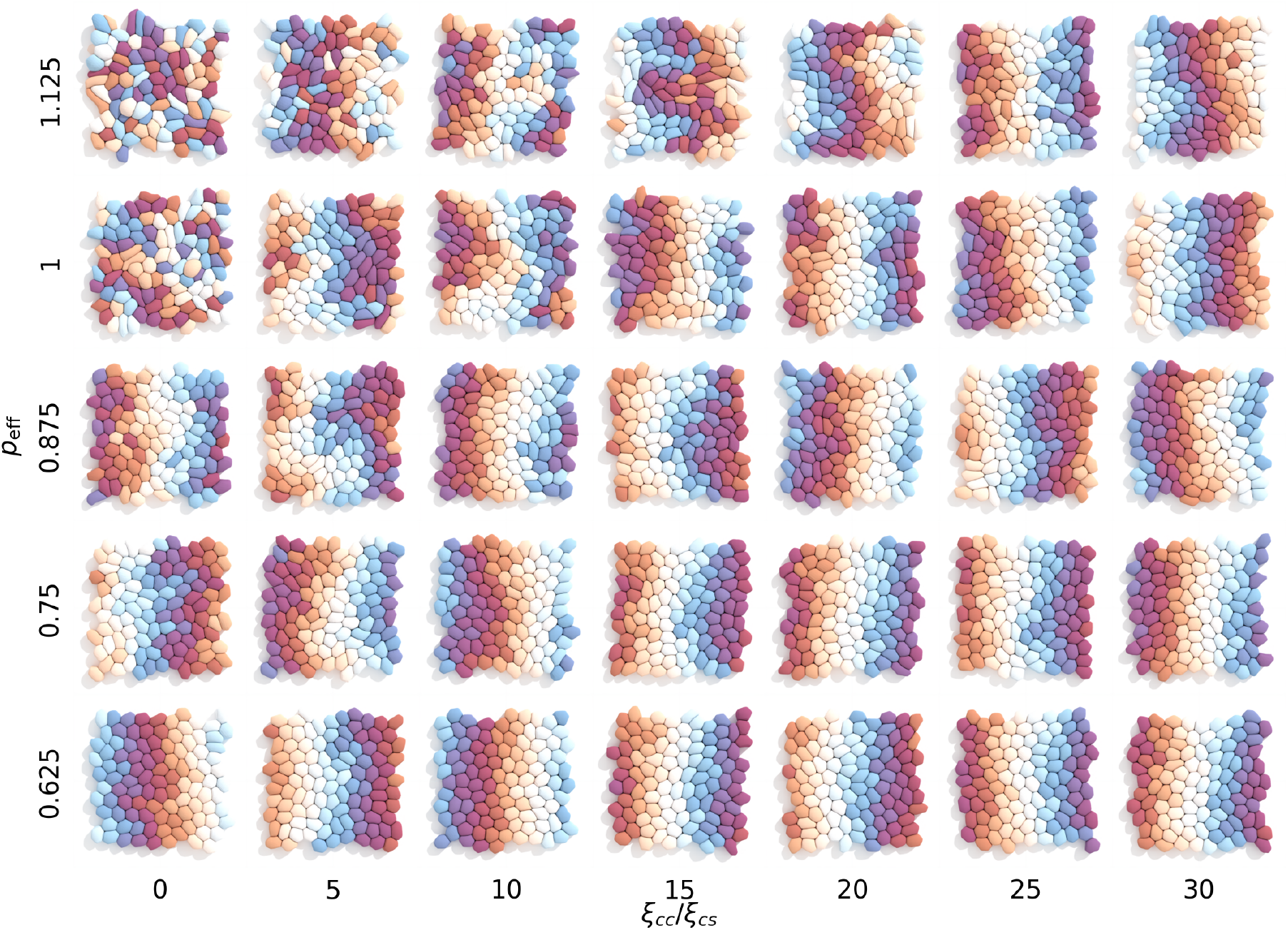
Snapshot of tissue at *t* = 50*τ*_*p*_ for confluent tissue (number density 1.1, *ω*_*cc*_*/γ* = 0.5), varying cell motility *p*_eff_ and intercellular friction *ξ*_*cc*_*/ξ*_*cs*_, where color indicates the initial position of the cell.

**Supplementary Fig. S8.**
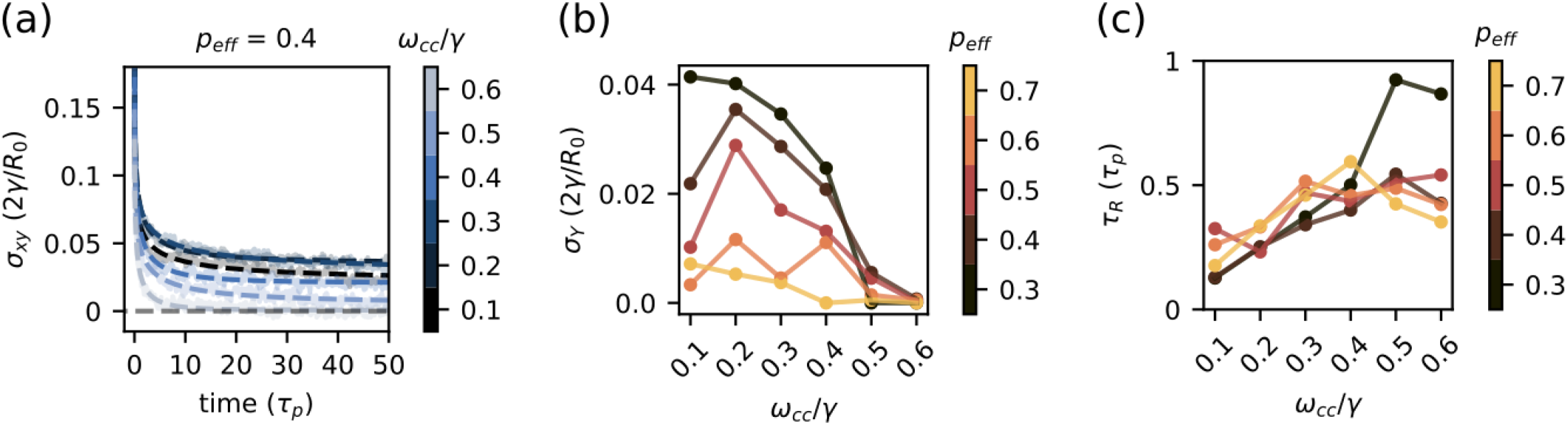
Shear stress relaxation for varying adhesive tension *ω*_*cc*_*/γ*, and cell activity *p*_eff_ at high number density (= 1.1). (a) Stretched exponential fit for *p*_eff_ = 0.4. (b) We show that yield stress *σ*_*Y*_ vanishes with increased activity, showing fluid tissue behavior, whereas adhesive tension has a non-monotonous effect on yield stress. At the high adhesive regime, *ω*_*cc*_*/γ* increases cell deformability, thereby lowering the energy barriers of T1-transitions, and ultimately steering the tissue towards fluid-like behavior. At the low adhesive regime, cell-separation forces increase with *ω*_*cc*_, therefore solidifying the tissue. (c) Relaxation time *τ*_*R*_ is primarily affected by *ω*_*cc*_*/γ* that increases effective viscosity of the tissue and *τ*_*R*_.

## References

[1] Elizabeth Lawson-Keister and M Lisa Manning. Jamming and arrest of cell motion in biological tissues. Current Opinion in Cell Biology, 72:146–155, 2021.

[2] Lior Atia, Jeffrey J Fredberg, Nir S Gov, and Adrian F Pegoraro. Are cell jamming and unjamming essential in tissue development? Cells & development, 168:203727, 2021.

[3] Yanlan Mao and Sara A Wickström. Mechanical state transitions in the regulation of tissue form and function. Nature Reviews Molecular Cell Biology, pages 1–17, 2024.

[4] Lior Atia, Dapeng Bi, Yasha Sharma, Jennifer A Mitchel, Bomi Gweon, Stephan A Koehler, Stephen J Decamp, Bo Lan, Jae Hun Kim, Rebecca Hirsch, Adrian F Pegoraro, Kyu Ha Lee, Jacqueline R Starr, David A Weitz, Adam C Martin, Jin ah Park, James P Butler, and Jeffrey J Fredberg. Geometric constraints during epithelial jamming. Nature Physics, 14:613–620, 2018.

[5] Alessandro Mongera, Payam Rowghanian, Hannah J. Gustafson, Elijah Shelton, David A. Kealhofer, Emmet K. Carn, Friedhelm Serwane, Adam A. Lucio, James Giammona, and Otger Campàs. A fluid-to-solid jamming transition underlies vertebrate body axis elongation. Nature, 561:401–405, 2018.

[6] Nicoletta I. Petridou, Silvia Grigolon, Guillaume Salbreux, Edouard Hannezo, and Carl Philipp Heisenberg. Fluidization-mediated tissue spreading by mitotic cell rounding and non-canonical wnt signalling. Nature Cell Biology, 21:169–178, 2019.

[7] Nicoletta I Petridou and Carl-Philipp Heisenberg. Tissue rheology in embryonic organization. The EMBO Journal, 38(20):e102497, 2019.

[8] Xun Wang, Matthias Merkel, Leo B Sutter, Gonca Erdemci-Tandogan, M Lisa Manning, and Karen E Kasza. Anisotropy links cell shapes to tissue flow during convergent extension. Proceedings of the National Academy of Sciences, 117:13541–13551, 2020.

[9] Nicoletta I Petridou, Bernat Corominas-Murtra, Carl-Philipp Heisenberg, and Edouard Hannezo. Rigidity percolation uncovers a structural basis for embryonic tissue phase transitions. Cell, 184(7):1914–1928, 2021.

[10] Sangwoo Kim, Rana Amini, Shuo-Ting Yen, Petr Pospíšil, Arthur Boutillon, Ilker Ali Deniz, and Otger Campàs. A nuclear jamming transition in vertebrate organogenesis. Nature Materials, 23(11):1592– 1599, 2024.

[11] Oleksandr Chepizhko, Maria Chiara Lionetti, Chiara Malinverno, Costanza Giampietro, Giorgio Scita, Stefano Zapperi, and Caterina AM La Porta. From jamming to collective cell migration through a boundary induced transition. Soft matter, 14(19):3774–3782, 2018.

[12] Robert J Tetley, Michael F Staddon, Davide Heller, Andreas Hoppe, Shiladitya Banerjee, and Yanlan Mao. Tissue fluidity promotes epithelial wound healing. Nature physics, 15(11):1195–1203, 2019.

[13] Jin Ah Park, Jae Hun Kim, Dapeng Bi, Jennifer A. Mitchel, Nader Taheri Qazvini, Kelan Tantisira, Chan Young Park, Maureen McGill, Sae Hoon Kim, Bomi Gweon, Jacob Notbohm, Robert Steward, Stephanie Burger, Scott H. Randell, Alvin T. Kho, Dhananjay T. Tambe, Corey Hardin, Stephanie A. Shore, Elliot Israel, David A. Weitz, Daniel J. Tschumperlin, Elizabeth P. Henske, Scott T. Weiss, M. Lisa Manning, James P. Butler, Jeffrey M. Drazen, and Jeffrey J. Fredberg. Unjamming and cell shape in the asthmatic airway epithelium. Nature Materials, 2015.

[14] Kenechukwu David Nnetu, Melanie Knorr, Josef Kas, and Mareike Zink. The impact of jamming on boundaries of collectively moving weak-interacting cells. New Journal of Physics, 14(11):115012, November 2012.

[15] Linda Oswald, Steffen Grosser, David M Smith, and Josef A Käs. Jamming transitions in cancer. Journal of physics D: Applied physics, 50(48):483001, 2017.

[16] Andrea Palamidessi, Chiara Malinverno, Emanuela Frittoli, Salvatore Corallino, Elisa Barbieri, Sara Sigismund, Galina V. Beznoussenko, Emanuele Martini, Massimiliano Garre, Ines Ferrara, Claudio Tripodo, Flora Ascione, Elisabetta A. Cavalcanti-Adam, Qingsen Li, Pier Paolo Di Fiore, Dario Parazzoli, Fabio Giavazzi, Roberto Cerbino, and Giorgio Scita. Unjamming overcomes kinetic and proliferation arrest in terminally differentiated cells and promotes collective motility of carcinoma. Nature Materials, pages 1–12, 2019.

[17] Jae Hun Kim, Adrian F. Pegoraro, Amit Das, Stephan A. Koehler, Sylvia Ann Ujwary, Bo Lan, Jennifer A. Mitchel, Lior Atia, Shijie He, Karin Wang, Dapeng Bi, Muhammad H. Zaman, Jin Ah Park, James P. Butler, Kyu Ha Lee, Jacqueline R. Starr, and Jeffrey J. Fredberg. Unjamming and collective migration in mcf10a breast cancer cell lines. Biochemical and Biophysical Research Communications, 521:706–715, 1 2020.

[18] Silvanus Alt, Poulami Ganguly, and Guillaume Salbreux. Vertex models: from cell mechanics to tissue morphogenesis. Philosophical Transactions of the Royal Society B: Biological Sciences, 372(1720):20150520, 2017.

[19] Dapeng Bi, J. H. Lopez, J. M. Schwarz, and M. Lisa Manning. A density-independent rigidity transition in biological tissues. Nature Physics, 11:1074–1079, 2015.

[20] Dapeng Bi, Xingbo Yang, M. Cristina Marchetti, and M. Lisa Manning. Motility-driven glass and jamming transitions in biological tissues. Physical Review X, 6:021011, 2016.

[21] Eial Teomy, David A Kessler, and Herbert Levine. Confluent and nonconfluent phases in a model of cell tissue. Physical Review E, 98(4):042418, 2018.

[22] Junxiang Huang, Herbert Levine, and Dapeng Bi. Bridging the gap between collective motility and epithelial– mesenchymal transitions through the active finite voronoi model. Soft Matter, 19(48):9389–9398, 2023.

[23] Sangwoo Kim, Marie Pochitaloff, Georgina Stooke-Vaughan, and Otger Campàs. Embryonic tissues as active foams. Nature Physics, 17:859–866, 2021.

[24] Hisao Honda, Masaharu Tanemura, and Tatsuzo Nagai. A three-dimensional vertex dynamics cell model of spacefilling polyhedra simulating cell behavior in a cell aggregate. Journal of theoretical biology, 226(4):439–453, 2004.

[25] Matthias Merkel and M Lisa Manning. A geometrically controlled rigidity transition in a model for confluent 3d tissues. New Journal of Physics, 20(2):022002, 2018.

[26] Tao Zhang and JM Schwarz. Boundary-bulk patterning in three-dimensional confluent cellular collectives. Physical Review Research, 43148:1–10, 2022.

[27] Elizabeth Lawson-Keister, Tao Zhang, Fatemeh Nazari, François Fagotto, and M Lisa Manning. Differences in boundary behavior in the 3d vertex and voronoi models. PLoS computational biology, 20(1):e1011724, 2024.

[28] Oliver M Drozdowski and Ulrich S Schwarz. Morphological instability at topological defects in a threedimensional vertex model for spherical epithelia. Physical Review Research, 6(2):L022045, 2024.

[29] Oliver M Drozdowski and Ulrich S Schwarz. Cell bulging and extrusion in a three-dimensional bubbly vertex model for curved epithelial sheets. arXiv preprint 2411.07141, 2024.

[30] Jean-Léon Mâitre, Héléne Berthoumieux, Simon Frederik Gabriel Krens, Guillaume Salbreux, Frank Jülicher, Ewa Paluch, and Carl-Philipp Heisenberg. Adhesion functions in cell sorting by mechanically coupling the cortices of adhering cells. Science, 338(6104):253–256, 2012.

[31] Rudolf Winklbauer. Cell adhesion strength from cortical tension–an integration of concepts. Journal of cell science, 128(20):3687–3693, 2015.

[32] Otger Campàs, Ivar Noordstra, and Alpha S. Yap. Adherens junctions as molecular regulators of emergent tissue mechanics. Nature Reviews Molecular Cell Biology, 4 2023.

[33] Ulrich S Schwarz and Samuel A Safran. Physics of adherent cells. Reviews of Modern Physics, 85(3):1327, 2013.

[34] Alexander Nestor-Bergmann, Guy B. Blanchard, Nathan Hervieux, Alexander G. Fletcher, Jocelyn Étienne, and Bénédicte Sanson. Adhesion-regulated junction slippage controls cell intercalation dynamics in an apposed-cortex adhesion model. PLOS Computational Biology, 18(1):1– 24, 01 2022.

[35] Michael Chiang, Austin Hopkins, Benjamin Loewe, M Cristina Marchetti, and Davide Marenduzzo. Intercellular friction and motility drive orientational order in cell monolayers. Proceedings of the National Academy of Sciences, 121(40):e2319310121, 2024.

[36] Alejandro Torres-Sánchez, Max Kerr Winter, and Guillaume Salbreux. Interacting active surfaces: A model for three-dimensional cell aggregates. PLoS Computational Biology, 18(12):e1010762, 2022.

[37] Christian Bächer, Diana Khoromskaia, Guillaume Salbreux, and Stephan Gekle. A three-dimensional numerical model of an active cell cortex in the viscous limit. Frontiers in Physics, 9:753230, 2021.

[38] Hudson Borja da Rocha, Jeremy Bleyer, and Hervé Turlier. A viscous active shell theory of the cell cortex. Journal of the Mechanics and Physics of Solids, 164:104876, 2022.

[39] Damien Cuvelier, Manuel Théry, Yeh-Shiu Chu, Sylvie Dufour, Jean-Paul Thiéry, Michel Bornens, Pierre Nassoy, and L Mahadevan. The universal dynamics of cell spreading. Current biology, 17(8):694–699, 2007.

[40] Elisabeth Fischer-Friedrich, Yusuke Toyoda, Cedric J Cattin, Daniel J Müller, Anthony A Hyman, and Frank Jülicher. Rheology of the active cell cortex in mitosis. Biophysical journal, 111(3):589–600, 2016.

[41] Arnab Saha, Masatoshi Nishikawa, Martin Behrndt, Carl-Philipp Heisenberg, Frank Jülicher, and Stephan W. Grill. Determining physical properties of the cell cortex. Biophysical Journal, 110(6):1421–1429, 2016.

[42] Sifan Yin, Bo Li, and Xi-Qiao Feng. Three-dimensional chiral morphodynamics of chemomechanical active shells. Proceedings of the National Academy of Sciences, 119(49):e2206159119, 2022.

[43] Hervé Turlier Basile Audoly, Jacques Prost, and Jean-François Joanny. Furrow constriction in animal cell cytokinesis. Biophysical journal, 106(1):114–123, 2014.

[44] Steve Runser, Roman Vetter, and Dagmar Iber. Simucell3d: three-dimensional simulation of tissue mechanics with cell polarization. Nature Computational Science, pages 1–11, 2024.

[45] Maxim Cuvelier, Jef Vangheel, Wim Thiels, Herman Ramon, Rob Jelier, and Bart Smeets. Stability of asymmetric cell division: A deformable cell model of cytokinesis applied to c. elegans. Biophysical Journal, 122(10):1858– 1867, 2023.

[46] Edouard Hannezo, Jacques Prost, and Jean Francois Joanny. Theory of epithelial sheet morphology in three dimensions. Proceedings of the National Academy of Sciences of the United States of America, 111:27–32, 2014.

[47] Françoise Brochard-Wyart and Pierre-Gilles de Gennes. Unbinding of adhesive vesicles. Comptes Rendus Physique, 4(2):281–287, 2003.

[48] Bart Smeets, Maxim Cuvelier, Jiri Pešek, and Herman Ramon. The effect of cortical elasticity and active tension on cell adhesion mechanics. Biophysical Journal, 116(5):930–937, 2019.

[49] Stéphane Douezan and Françoise Brochard-Wyart. Dewetting of cellular monolayers. The European Physical Journal E, 35:1–6, 2012.

[50] Ivana Pajic-Lijakovic, Milan Milivojevic, and Peter V. E. McClintock. Epithelial cell-cell interactions in an overcrowded environment: jamming or live cell extrusion. Journal of Biological Engineering, 18(1), September 2024.

[51] Eva Maria Schötz, Marcos Lanio, Jared A. Talbot, and M. Lisa Manning. Glassy dynamics in three-dimensional embryonic tissues. Journal of the Royal Society Interface, 10:20130726, 2013.

[52] Simon Garcia, Edouard Hannezo, Jens Elgeti, Jean-François Joanny, Pascal Silberzan, and Nir S. Gov. Physics of active jamming during collective cellular motion in a monolayer. Proceedings of the National Academy of Sciences, 112(50):15314–15319, 2015.

[53] A. J. Liu and S. R. Nagel. Jamming is not just cool any more. Nature, 396:21–22, 1998.

[54] V. Trappe, V. Prasad, Luca Cipelletti, P.N. Segre, and D.A. Weitz. Jamming phase diagram for attractive particles. Nature, 411:772–775, 2001.

[55] Xun Wang, Christian M. Cupo, Sassan Ostvar, Andrew D. Countryman, and Karen E. Kasza. E-cadherin tunes tissue mechanical behavior before and during morphogenetic tissue flows. Current Biology, 34(15):3367– 3379.e5, 2024.

[56] Kevin K. Chiou, Lars Hufnagel, and Boris I. Shraiman. Mechanical stress inference for two dimensional cell arrays. PLOS Computational Biology, 8(5):1–9, 05 2012.

[57] K. Venkatesan Iyer, Romina Piscitello-Gómez, Joris Paijmans, Frank Jülicher, and Suzanne Eaton. Epithelial viscoelasticity is regulated by mechanosensitive e-cadherin turnover. Current Biology, 29(4):578–591.e5, 2019.

[58] Jin-Ah Park, Lior Atia, Jennifer A. Mitchel, Jeffrey J. Fredberg, and James P. Butler. Collective migration and cell jamming in asthma, cancer and development. Journal of Cell Science, 129(18):3375–3383, 09 2016.

[59] Marielena Gamboa Castro, Susan E. Leggett, and Ian Y. Wong. Clustering and jamming in epithelial–mesenchymal co-cultures. Soft Matter, 12:8327–8337, 2016.

[60] Sophie Lohmann, Costanza Giampietro, Francesca M. Pramotton, Dunja Al-Nuaimi, Alessandro Poli, Paolo Maiuri, Dimos Poulikakos, and Aldo Ferrari. The role of tricellulin in epithelial jamming and unjamming via segmentation of tricellular junctions. Advanced Science, 7, 8 2020.

[61] Ian T. Stancil, Jacob E. Michalski, Duncan Davis-Hall, Hong Wei Chu, Jin Ah Park, Chelsea M. Magin, Ivana V. Yang, Bradford J. Smith, Evgenia Dobrinskikh, and David A. Schwartz. Pulmonary fibrosis distal airway epithelia are dynamically and structurally dysfunctional. Nature Communications, 12, 12 2021.

[62] Jennifer A. Mitchel, Amit Das, Michael J. O’Sullivan, Ian T. Stancil, Stephen J. DeCamp, Stephan Koehler, Oscar H. Ocaña, James P. Butler, Jeffrey J. Fredberg, M. Angela Nieto, Dapeng Bi, and Jin Ah Park. In primary airway epithelial cells, the unjamming transition is distinct from the epithelial-to-mesenchymal transition. Nature Communications, 11:1–14, 2020.

[63] Grace Cai, Nicole C. Rodgers, and Allen P. Liu. Unjamming transition as a paradigm for biomechanical control of cancer metastasis, 2024.

[64] Steve Pawlizak, Anatol W. Fritsch, Steffen Grosser, Dave Ahrens, Tobias Thalheim, Stefanie Riedel, Tobias R. Kießling, Linda Oswald, Mareike Zink, M. Lisa Manning, and Josef A. Käs. Testing the differential adhesion hypothesis across the epithelial-mesenchymal transition. New Journal of Physics, 17:083049, 2015.

[65] Julia Eckert, Benòit Ladoux, René Marc Mège, Luca Giomi, and Thomas Schmidt. Hexanematic crossover in epithelial monolayers depends on cell adhesion and cell density. Nature Communications, 14, 12 2023.

[66] Monirosadat Sadati, Nader Taheri Qazvini, Ramaswamy Krishnan, Chan Young Park, and Jeffrey J. Fredberg. Collective migration and cell jamming. Differentiation, 86(3):121–125, 2013. Mechanotransduction.

[67] L Gay, AM Baker, and TA Graham. Tumour cell heterogeneity [version 1; peer review: 5 approved]. F1000Research, 5(238), 2016.

[68] Hervé Turlier and Jean-Léon Mâitre. Mechanics of tissue compaction. Seminars in Cell Developmental Biology, 47-48:110–117, 2015. Coding and non-coding RNAs Mammalian development.

[69] Martin Bergert, Anna Erzberger, Ravi A Desai, Irene M Aspalter, Andrew C Oates, Guillaume Charras, Guillaume Salbreux, and Ewa K Paluch. Force transmission during adhesion-independent migration. Nature cell biology, 17(4):524–529, 2015.

[70] AC Callan-Jones, J-F Joanny, and J Prost. Viscousfingering-like instability of cell fragments. Physical review letters, 100(25):258106, 2008.

[71] Kelly Vazquez, Aashrith Saraswathibhatla, and Jacob Notbohm. Effect of substrate stiffness on friction in collective cell migration. Scientific Reports, 12(1):2474, 2022.

[72] Yuansheng Cao, Richa Karmakar, Elisabeth Ghabache, Edgar Gutierrez, Yanxiang Zhao, Alex Groisman, Herbert Levine, Brian A. Camley, and Wouter-Jan Rappel. Cell motility dependence on adhesive wetting. Soft Matter, 15:2043–2050, 2019.

[73] Steven Ongenae, Hanna Svitina, Tom E. R. Belpaire, Jef Vangheel, Tobie Martens, Pieter Vanden Berghe, Ioannis Papantoniou, and Bart Smeets. Active foam dynamics of tissue spheroid fusion. bioRxiv, 2024.

[74] D. Fedosov, B. Caswell, and G. Karniadakis. Systematic coarse-graining of spectrin-level red blood cell models. Computater Methods Applied Mechanics and Engineering, 199:29–32, 2010.

[75] Bart Smeets, Tim Odenthal, Simon Vanmaercke, and Herman Ramon. Polygon-based contact description for modeling arbitrary polyhedra in the discrete element method. Computer Methods in Applied Mechanics and Engineering, 290:277–289, 2015.

[76] Kenneth A. Brakke. The surface evolver. Experimental Mathematics, 1:141–165, 1992.

[77] J. H. Irving and John G. Kirkwood. The statistical mechanical theory of transport processes. iv. the equations of hydrodynamics. The Journal of Chemical Physics, 18(6):817–829, 06 1950.

[78] Paul Liedekerke, Margriet Palm, Nick Jagiella, and Dirk Drasdo. Simulating tissue mechanics with agent-based models: concepts, perspectives and some novel results. Computational Particle Mechanics, 2:401–444, 2015.

[79] https://gitlab.kuleuven.be/mebios-particulate/publications/rigidity_transitions_cell_ monolayers.git.

