## Supplementary Material for "Rigidity transitions in a 3D active foam model of cell monolayers with frictional contact interactions"

April 17, 2025

#### Theoretical considerations on foam mechanics

##### Equilibrium shape of cells in a hexagonal confluent tissue

Assume that a cell with cortex tension $\gamma$ spreads out on a substrate with adhesive tension $\omega_{cs}$ and establishes a hexagonal confluent tissue with adhesive tension $\omega_{cc}$ between neighboring cells. Assuming volume conservation, the free energy of the cell can be written as

$$\mathscr{F} = \gamma S_a + (\gamma - \omega_{cc}) S_l + (\gamma - \omega_{cs}) S_b$$

where $S_a = S_b$ is the apical and basal cell surface area and $S_l$ is the cell area in contact with neighboring cells. Let us define $h$ as the cell height and $d$ as the distance between the centroids of neighboring cells to rewrite the free energy as

$$\mathscr{F} = (2\gamma - \omega_{cs}) \frac{\sqrt{3}}{2} d^2 + (\gamma - \omega_{cc}) \frac{6}{\sqrt{3}} d\,h.$$

Given the cell is initially spherical with radius $R_0$, and assuming volume conservation ($V_0 = 4/3\pi R_0^3$), we can write cell height $h$ in function of cell-cell distance $d$ as

$$h = \frac{8\pi R_0^3}{3\sqrt{3} d^2}$$

$$d = \sqrt{\frac{8\pi R_0^3}{3\sqrt{3} h}}$$

We use this relation to write the free energy in function of cell-cell centroid distance

$$\mathscr{F} = (2\gamma - \omega_{cs}) \frac{\sqrt{3}}{2} d^2 + (\gamma - \omega_{cc}) \frac{16\pi}{3} \frac{R_0^2}{d}$$

or in function of cell height

$$\mathscr{F} = (2\gamma - \omega_{cs}) \sqrt{V_0} h + (\gamma - \omega_{cc}) \sqrt{V_0 8\sqrt{3} h}.$$

By minimizing the free energy $\frac{\mathrm{d}\mathscr{F}}{\mathrm{d}d} = 0$, we can calculate the cell shape corresponding to minimal energy in function of cortex tension and adhesive tensions. The derivative of free energy in function of $d$

$$\frac{\mathrm{d}\mathscr{F}}{\mathrm{d}d} = (2\gamma - \omega_{cs}) \sqrt{3} d - (\gamma - \omega_{cc}) \frac{16\pi R_0^3}{3d^2},$$

so we find the corresponding cell-cell distance:

$$d/R_0 = \sqrt[3]{\frac{16\pi}{3\sqrt{3}}\left(\frac{\gamma-\omega_{cc}}{2\gamma-\omega_{cs}}\right)} \tag{S1}$$

and cell height

$$h/R_0 = \left(\frac{2\pi}{3\sqrt{3}}\right)^{1/3}\left(\frac{\gamma-\omega_{cc}}{2\gamma-\omega_{cs}}\right)^{-2/3}$$

From this, we calculate the ratio

$$h/d = \frac{1}{2}\left(\frac{2\gamma-\omega_{cs}}{\gamma-\omega_{cc}}\right). \tag{S2}$$

As expected, this shows that cell-substrate adhesive tension $\omega_{cs}$ promotes squamous cell shapes whereas cell-substrate adhesive tension $\omega_{cc}$ promotes columnar shapes. Lastly, we may also find the apical area of the cell

$$S_a = \frac{\sqrt{3}}{2}R_0^2\left(\frac{16\pi}{3\sqrt{3}}\frac{\gamma-\omega_{cc}}{2\gamma-\omega_{cs}}\right)^{2/3}$$

#### Active dipole model of cell migration

Cell activity is captured by two distinct but related effects: a net protrusive force and a force dipole. The net traction force of the cell is balanced by friction forces to drive cell movement at a constant velocity $v_0$ (Fig. 1 a). Furthermore, an active dipole $\nabla_p$ stretches the cell in the direction of polarization $\mathbf{p}$ (Fig. 1 b). The pressure on the cell surface at position $\mathbf{x}_i$ from cell activity is given by

$$P_a(\mathbf{x}_i) = \left(0.5p_a\frac{|(\mathbf{x}-\mathbf{x}_0)\cdot\mathbf{p}|}{L_c} + \nabla_p(\mathbf{x}_i-\mathbf{x}_0)\cdot\mathbf{p}\right)(\mathbf{n}_i\cdot\mathbf{p})$$

where $p_a = v_0\xi_s$ with $\xi_s$ the cell-substrate friction, and $\mathbf{n}_i$ is the normal direction of the cell surface at position $\mathbf{x}_i$. We assume both $v_0$ and $\nabla_p$ scale by the same active parameter $p_a$ (Fig. 1 c):

$$P_a(\mathbf{x}_i) = p_a\left(0.5\frac{|(\mathbf{x}-\mathbf{x}_0)\cdot\mathbf{p}|}{L_c} + \frac{(\mathbf{x}_i-\mathbf{x}_0)}{\lambda_a L_c}\cdot\mathbf{p}\right)(\mathbf{n}_i\cdot\mathbf{p})$$

such that $\nabla_p = p_a/(\lambda_a L_c)$ where $L_c$ is the instantaneous length of the cell in the direction of polarization and $\lambda_a$ is the dipole scaling factor: smaller implies a stronger dipole. Furthermore, we assume the cell migrates by actin polymerization close to the substrate, so we limit the protrusive forces to a narrow region near the substrate (located parallel to the $xz$ plane at position $y = y_{sub}$) by multiplying $P_a(\mathbf{x}_i)$ with a Gaussian kernel with a length scale $\lambda_z$ similar to [1]:

$$P_a(\mathbf{x}_i) = p_a\left(0.5\frac{|(\mathbf{x}-\mathbf{x}_0)\cdot\mathbf{p}|}{L_c} + \frac{(\mathbf{x}_i-\mathbf{x}_0)}{\lambda_a L_c}\cdot\mathbf{p}\right)(\mathbf{n}_i\cdot\mathbf{p})\exp\left[-\frac{1}{2}\left(\frac{y_i-y_{sub}}{\lambda_z^2}\right)^2\right] \tag{S3}$$

##### Cell elongation from active force dipole

The force dipole $\nabla_p$ works in a similar way as the preferred shape index $q_0$ in vertex models. Let's observe the cell from the basal side in a 2D perspective and assume that the basal area deforms from a circular to an elliptical shape with semi-major axis $a$ and semi-minor axis $b$. The cell periphery is given by $[a\sin(\phi), b\cos(\phi)]$ such that the radius of curvature of an ellipse can be written as

$$r(\phi) = \frac{(a^2\sin^2(\phi)+b^2\cos^2(\phi))^{3/2}}{ab}$$

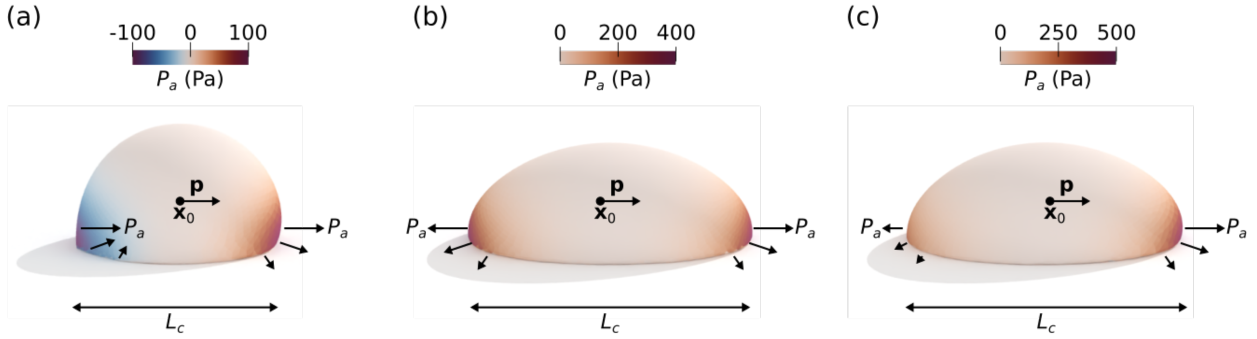


Fig. 1: Example of a single migrating cell (a) with only migratory forces $p_a = 200$ Pa and no active dipole $\nabla_p = 0$ Pa m$^{-1}$, (b) with no migratory forces $p_a = 0$ Pa but only an active dipole $\nabla_p = 400$ Pa m$^{-1}$, and (c) both migratory forces and active dipole as given in eq. S3.

From the Young-Laplace law $\Delta P = 2\gamma_{\text{eff}}/r$, we can calculate the expected curvature in the direction of the dipole $\phi = 0$ and orthogonal to the dipole $\phi = \pi/2$

$$\Delta P = \begin{cases} \frac{2\gamma_{\text{eff}}}{a^2/b}, & \phi = 0 \\ \frac{2\gamma_{\text{eff}}}{b^2/a} - \frac{p_a}{\lambda_a}, & \phi = \pi/2 \end{cases}$$

that can be rewritten as

$$\frac{p_a}{\lambda_a} = 2\gamma_{\text{eff}}\left(\frac{a}{b^2} - \frac{b}{a^2}\right).$$

Assuming the basal area of the cell is conserved ($S_a = \pi ab = \pi R_{2D}^2$) we can find the relation between $a$ and $b$. From this, we rewrite the balance as

$$\frac{p_a}{\lambda_a} = 2\gamma_{\text{eff}}\left(\frac{a^3}{R_{2D}^4} - \frac{R_{2D}^2}{a^3}\right).$$

or

$$\frac{p_a R_{2D}}{2\gamma_{\text{eff}}\lambda_a} = \frac{a^3}{R_{2D}^3} - \frac{R_{2D}^3}{a^3}.$$

$$\frac{p_a R_{2D}}{2\gamma_{\text{eff}}\lambda_a} = \frac{a^6 - R_{2D}^6}{a^3 R_{2D}^3}$$

We can solve $a$ and $b$ in function of the non-dimensionalized active dipole pressure $p_{0,\text{eff}} = \frac{p_a R_{2D}}{2\gamma_{\text{eff}}\lambda_a}$:

$$a = R_{2D}\left(\frac{p_{0,\text{eff}} + \sqrt{p_{0,\text{eff}}^2 + 4}}{2}\right)^{1/3}$$

$$b = R_{2D}\left(\frac{p_{0,\text{eff}} + \sqrt{p_{0,\text{eff}}^2 + 4}}{2}\right)^{-1/3}$$

The aspect ratio $\mathscr{A} = a/b$ of the basal area of the cell becomes

$$\frac{a}{b} = \left(\frac{p_{0,\text{eff}}}{2} + \sqrt{\frac{p_{0,\text{eff}}{}^2}{4} + 1}\right)^{2/3}$$

Solving to $p_{0,\text{eff}}$ gives

$$p_{0,\text{eff}} = \frac{\mathscr{A}^3 - 1}{\mathscr{A}^{3/2}}$$

According to vertex theory, the tissue should unjam when a critical perimeter-to-surface ratio or aspect ratio is reached $\mathscr{A}^*$. Assuming $\gamma_{\text{eff}} = \gamma - \omega_{cc}$ we can calculate the critical active dipole $p_0^*$ where $p_0 = \frac{p_a R_{2D}}{2\gamma\lambda_a}$ as

$$p_0^* = \frac{\mathscr{A}^{*3} - 1}{\mathscr{A}^{*3/2}}(1 - \omega_{cc}/\gamma)$$

Finally, we can use the calculations in eq. S1 to find a relation between $R_{2D}$ and $\gamma_{\text{eff}} = \gamma - \omega_{cc}$

$$R_{2D} \propto d/2 \propto R_0 \left(\frac{\gamma - \omega_{cc}}{2\gamma - \omega_{cs}}\right)^{1/3}$$

so that the critical activity $p^*_{\text{eff}} = \left(\frac{p_a R_0}{2\gamma\lambda}\right)^*$ is given by

$$p^*_{\text{eff}} \propto \frac{\mathscr{A}^{*3} - 1}{\mathscr{A}^{*3/2}}(1 - \omega_{cc}/\gamma)^{2/3}(2 - \omega_{cs}/\gamma)^{1/3} \tag{S4}$$

where we see that $\omega_{cc}$ and $\omega_{cs}$ will promote cell deformation, and according to vertex theory, tissue unjamming. Where $\omega_{cc}$ decreases cell interfacial tension that counteracts deformation, $\omega_{cs}$ promotes deformation by increasing the apical area $\pi R_{2D}^2$ of the cell thereby reducing the pressure needed for cell deformation (from Young-Laplace law $\Delta P = 2\gamma/r$).

**Cell-cell pulloff**

In the limit of porous tissue, cell-cell pulloff is the main driving factor of cell neighbor exchanges and thus tissue unjamming. The two opposing forces determining tissue cohesion is cell-cell rupture force, and the active pulling force. The rupture force, previously derived by Brochart-Wyart and De Gennes [2], is approximated by

$$F_s \approx \pi R_0 \omega_{cc}.$$

The pulling force results from active migration of cells and is approximated by

$$F_a \approx p_a R_0 \lambda_z.$$

The tissue cohesion parameter $p_a^*$ becomes

$$\pi R_0 \omega_{cc} = p_a^* R_0 \lambda_z$$

or

$$p_a^* \propto \frac{\pi \omega_{cc}}{\lambda_z}. \tag{S5}$$

At low cell-cell adhesive tension and cell density, we expect the critical protrusive pressure $p_a^*$, where tissue changes from jammed to unjammed, to be proportional to $\omega_{cc}$.

### Numerical implementation of active foam model

In this section we provide the implementation of the discretized active foam model previously described by [3, 4]. In this model, cells are represented as pressurized bubbles with a viscous cortex under tension and adhesive bonds allowing cell-cell and cell-substrate contact. The cell actomyosin complex is represented by a triangulated surface mesh where the vertex positions $\mathbf{x}_i$ act as the relevant degrees of freedom.

#### Cortex tension

Tension in the cortex arises from actomyosin contractility, modeled as an effective surface tension $\gamma_c$. At contact interfaces, cortex tension decreases by decrease of actomyosin activity [5] represented as an effective adhesive tension $\omega_{cj}$. The resulting surface tension in a triangle $A$ is given by

$$\gamma_c = \begin{cases} \gamma - \omega_{cj}, & \text{for contacting surfaces and,} \\ \gamma & \text{for free surfaces.} \end{cases}$$

The force on each node $i$ of a triangle consisting out of nodes $A = (i, j, k)$ is then given by [6]

$$\mathbf{F}_i^{act} = \frac{\gamma_c}{2}(\mathbf{x}_k - \mathbf{x}_j) \times \hat{\mathbf{n}}_A$$

#### Cortex viscosity

Actomyosin remodeling is modeled as an effective viscosity $\eta_c$. We distinguish between in-plane and out-of-plane bending viscosity. The in-plane viscous damping force between two nodes $i, j$ with velocity $\mathbf{u}$ is given by

$$\mathbf{F_i}^{\text{visc}} = \frac{\eta_c t_c}{\sqrt{3}} \left( (\hat{\mathbf{n}}_{ij} \cdot (\mathbf{u}_j - \mathbf{u}_i))\hat{\mathbf{n}}_{ij} + (\hat{\mathbf{t}}_{ij} \cdot (\mathbf{u}_j - \mathbf{u}_i))\hat{\mathbf{t}}_{ij} \right),$$

where the direction of viscous forces are given by the unit vectors

$$\hat{\mathbf{n}}_{\mathbf{ij}} = \frac{\mathbf{x_j} - \mathbf{x_i}}{||\mathbf{x_j} - \mathbf{x_i}||},$$

$$\hat{\mathbf{t}}_{\mathbf{ij}} = \frac{\hat{\mathbf{n}}_{\mathbf{A}} + \hat{\mathbf{n}}_{\mathbf{B}}}{||\hat{\mathbf{n}}_{\mathbf{A}} + \hat{\mathbf{n}}_{\mathbf{B}}||} \times \hat{\mathbf{n}}_{\mathbf{ij}},$$

with $\hat{\mathbf{n}}_{\mathbf{A}}$, and $\hat{\mathbf{n}}_{\mathbf{B}}$, the normals of the triangles making up the segment between nodes $i$ and $j$. Note that we do not distinguish between bulk, extensional ($\eta_e$) or shear viscosity ($\eta_c$), effectively setting the Trouton's ratio to $\eta_e/\eta_c = 1$, whereas a Newtonian fluid would have a value equal 3.

The viscous bending moment between two adjacent triangles $A = (i, j, k), B = (i, l, j)$ with small thickness $t_c$ arises from the rate of angle deviation $\frac{\mathrm{d}\phi}{\mathrm{dt}}$ as

$$||\mathbf{M}_{AB}^{\text{visc}}|| = \frac{\eta_c t_c^3}{3} \frac{\mathrm{d}\phi}{\mathrm{dt}}$$

From this bending moment, we compute the lever forces on the opposing nodes

$$\mathbf{F}_k^{\text{visc}} = -\frac{\mathbf{M}_{\mathbf{AB}}^{\text{visc}} \times \hat{\mathbf{l}}_A}{l_A},$$

$$\mathbf{F}_l^{\text{visc}} = +\frac{\mathbf{M}_{\mathbf{AB}}^{\text{visc}} \times \hat{\mathbf{l}}_B}{l_B},$$

with $l$ and $\hat{\mathbf{l}}$ the length and unit vector from common axis $(i, j)$ to lever point $k$ or $l$. For lever $A$ it is computed as

$$\mathbf{l_A} = \hat{\mathbf{n}}_A \times (\mathbf{x}_j - \mathbf{x}_i),$$

$$l_A = (\mathbf{x}_k - \mathbf{x}_i) \cdot \mathbf{l_A}.$$

Analogously, we can calculate it for lever $B$. The forces on the common points $(i, j)$ are weighted with the projected distance of the lever on the common axis

$$y_{i,A} = (\mathbf{x}_k - \mathbf{x}_i) \cdot (\mathbf{x}_j - \mathbf{x}_i)$$

$$y_{j,A} = (\mathbf{x}_j - \mathbf{x}_k) \cdot (\mathbf{x}_j - \mathbf{x}_i)$$

$$y_{i,B} = (\mathbf{x}_l - \mathbf{x}_i) \cdot (\mathbf{x}_j - \mathbf{x}_i)$$

$$y_{j,B} = (\mathbf{x}_j - \mathbf{x}_l) \cdot (\mathbf{x}_j - \mathbf{x}_i)$$

to obtain the forces on the common nodes for lever forces $F_k^{\text{visc}}$ and $F_l^{\text{visc}}$

$$\mathbf{F}_i^{\text{visc}} = -\frac{y_{j,A}}{y_{i,A} + y_{j,A}} \mathbf{F}_k^{\text{visc}} - \frac{y_{j,B}}{y_{i,B} + y_{j,B}} \mathbf{F}_l^{\text{visc}},$$

$$\mathbf{F}_j^{\text{visc}} = -\frac{y_{i,A}}{y_{j,A} + y_{i,A}} \mathbf{F}_k^{\text{visc}} - \frac{y_{i,B}}{y_{i,B} + y_{j,B}} \mathbf{F}_l^{\text{visc}},$$

that ensures force balance.

#### Internal pressure

We assume the cell cytoplasm to be incompressible. Therefore, we implement active volume control for each cell by a cytoplasmic pressure $P_b$ using a PI volume-controller

$$P_b = -K\varepsilon_V(t) - K_i \int_0^t \varepsilon_V(t)\mathrm{d}t,$$

where $\varepsilon_V(t) = (V(t) - V_0)/V_0$, with $V(t)$ the instantaneous volume and $V_0$ the equilibrium volume, $K$ the proportional and $K_i$ integrative term. The force on a triangle $A$ is given by

$$\mathbf{F}_A^b = \mathbf{S}_A P_b(t),$$

with $\mathbf{S}_A$ the directed area of the triangle, that is simply distributed to the nodes $i$ as

$$\mathbf{F}_i^b = \frac{\mathbf{F}_A^b}{3}.$$

#### Active migration pressure

Cell migration is represented by protrusive forces at the cell front end and retraction forces at the trailing end. Typically, cells elongate during migration, modeled as a dipole pressure. The pressure at triangles $A$ is given by

$$P_A^a(\mathbf{x}) = \left(\frac{p_a}{2} + \nabla_p(\mathbf{x} - \mathbf{x}_0) \cdot \hat{\mathbf{p}})(\hat{\mathbf{n}}_A \cdot \hat{\mathbf{p}}\right) \exp\left[-\frac{1}{2}\left(\frac{z - z_s}{\lambda_z^2}\right)^2\right],$$

where $\mathbf{x}$ is the geometric center of the triangle, $\hat{\mathbf{n}}_A$ the triangle normal, $\mathbf{x}_0$ is the geometric center of the cell, and $\hat{\mathbf{p}}$ the cell polarization vector. The triangle forces $\mathbf{F}_A^a = \mathbf{S}_A P_A^a(t)$ are distributed to the nodes $i$ as

$$\mathbf{F}_i^a = \frac{\mathbf{F}_A^a}{3}.$$

#### Contact pressure

We assume uniform adhesion over interacting surfaces. The contact pressure $P^c_{AB}$ includes repulsive and adhesive interactions as

$$P^c_{AB}(\mathbf{x}) = k\delta(\mathbf{x}) - P^0$$

where $P^0 = \omega/h_0$, with $h_0$ the effective range of adhesion that is accomplished by translating the nodes over this distance $h_0$ in the direction of the node normals. To ensure that at equilibrium the work of adhesion is recovered, the stiffness is set as $k = P^0/h_0$. The contact overlap at point $\mathbf{x}$ on a triangle is computed with the translated triangles as

$$\delta(\mathbf{x}) = \max\left(0; 2(\mathbf{x} - \mathbf{s}_{AB}) \cdot (\hat{\mathbf{n}}_{AB} \times \mathbf{I}_{AB}) \tan(\alpha)\right)$$

with $\mathbf{I}_{AB}$ the intersection line between two triangles, $\mathbf{s}_{AB}$ an arbitrary point on the intersection line, $\hat{\mathbf{n}}_{AB}$ the normal of the common contact plane defined as

$$\hat{\mathbf{n}}_{AB} = \frac{\hat{\mathbf{n}}_A - \hat{\mathbf{n}}_B}{\|\hat{\mathbf{n}}_A - \hat{\mathbf{n}}_B\|},$$

and $\alpha$ the angle between the triangle and the contact plane $A \cap B$. The contact force and moment is obtained by integrating the total contact pressure over the oriented contact area $\mathbf{S}_{AB}$ on the common contact plane

$$\mathbf{F}^c_{AB} = \int_{\mathbf{x} \in A \cap B} P^c_{AB}(\mathbf{x}) d\mathbf{S}_{AB},$$

$$\mathbf{M}^c_{AB} = \int_{\mathbf{x} \in A \cap B} P^c_{AB}(\mathbf{x}) d\mathbf{S}_{AB} \times (\mathbf{x} - \mathbf{x}_{AB}),$$

with $\mathbf{x}_{AB}$ the contact point defined as the geometric center of the intersection polygon. This force is distributed to the nodes $i$ ensuring conservation of momentum and force, assuming the forces are directed in $\hat{\mathbf{n}}_{AB}$. This results in a system of linear equations per contact pair

$$\mathbf{M}^c_{AB} = \sum_{i \in A} (\mathbf{x} - \mathbf{x}_{AB}) \times \mathbf{F}^c_{i,AB},$$

$$\mathbf{F}^c_{AB} = \sum_{i \in A} \mathbf{F}^c_{i,AB}.$$

The solution for this system is presented and discussed in the work of Smeets et al. [7].

#### Wet contact friction

A viscous contact force is included to account for drag between contacting surfaces with friction constant $\xi_{ci}$. The contact drag force acting on node $i$ of triangle $A$ of the contact pair $(AB)$ is computed as

$$\mathbf{F}^{\text{fric}}_{AB,i} = \Gamma^{\text{fric}}_{AB} \cdot \sum_{k \in B} w^{AB}_{ik} \left(\mathbf{u}_k - \mathbf{u}_i\right)$$

determined by a friction tensor $\Gamma^{\text{fric}}_{AB}$ and weights $w^{AB}_{ik}$ per node $k$ of the $B$ triangle. $w^{AB}_{ik}$ are assumed to scale with the relative contribution of the nodal contact forces to the overall contact force thus

$$w^{AB}_{ik} = \frac{\left(\mathbf{F}^c_{AB,i} + \mathbf{F}^c_{AB,k}\right) \cdot \hat{\mathbf{n}}_{AB}}{6 \sum_{\forall k \in B} \mathbf{F}^c_{AB,k} \cdot \hat{\mathbf{n}}_{AB}}$$

is used as an approximation. $\Gamma^{\text{fric}}_{AB}$ for a given contact area $S_{AB}$ is estimated as

$$\Gamma^{\text{fric}}_{AB} = S_{AB} \left[\xi^{\perp} \hat{\mathbf{n}}_{AB} \otimes \hat{\mathbf{n}}_{AB} + \xi^{\parallel} \left(-\hat{\mathbf{n}}_{AB} \otimes \hat{\mathbf{n}}_{AB}\right)\right],$$

with normal and tangential friction coefficients $\xi^{\perp}$ and $\xi^{\parallel}$ respectively.

#### Medium damping

A general drag force on node $i$ from liquid drag between cell and medium with viscosity $\eta_f$ is given by

$$\mathbf{F}_i^{\text{drag}} = -\Gamma_i^{\text{f}} \cdot \mathbf{u}_i,$$

where we approximate the drag force from Stokes' Law for spherical particles as

$$\Gamma_i^{\text{f}} = 6\pi R_0 \eta_f \frac{S_i}{4\pi R_0^2} = \frac{3\eta_f}{2R_0} S_i.$$

$F_i^{\text{drag}}$ is small compared to other dissipative forces, but provides stability of the numerical integration scheme as it ensures the resistance matrix is positive definite.

#### Equation of motion

For the overdamped environment, interial forces can be neglected and the force balance for node $i$ is given by

$$\mathbf{F}_i^{\text{act}} + \mathbf{F}_i^{\text{b}} + \mathbf{F}_i^{\text{a}} + \sum_{i \in A} \mathbf{F}_{i,AB}^{c} = \mathbf{F}_i^{\text{visc}} + \sum_{i \in A} \mathbf{F}_{i,AB}^{\text{fric}} + \mathbf{F}_i^{\text{drag}},$$

where right-hand forces are viscous forces that depend on relative velocities of the nodes. For the total system of $N$ nodes, this can be written as

$$\mathbf{\Lambda} \mathbf{u} = \mathbf{F}, \tag{S6}$$

with $\mathbf{F}$ a $3N \times 1$ column matrix that represents all left-hand forces per node, $\mathbf{\Lambda}$ a $3N \times 3N$ sparse symmetric positive definite friction matrix, and $\mathbf{u}$ a $3N \times 1$ column matrix representing the velocities of $N$ nodes. The congujate gradient method (CGM) is then used to solve for the node velocities every timestep. Node positions $\mathbf{x}_i$ are updated every timestep using a semi-implicit Euler integration scheme

$$\mathbf{x}_i(t + \Delta t) = \mathbf{x}_i(t) + \Delta t\, \mathbf{u}_i(t + \Delta t),$$

where $\mathbf{u}_i(t + \Delta t)$ are the projected new velocities obtained by solving eq. S6 at time $t$ via the CGM.

#### Remeshing

Because we model the cortex as a viscous fluid under surface tension, the discrete nodes tend to flow over the surface. To ensure the mesh quality, we remesh the triangulated surface. There are 4 remeshing stages based on [8]: split triangles, flip edges, split edges, and delete edges. In the first stage, triangles that are too big ($S_A > S_{\min}$) are split at the geometric center in 3 separate triangles. In the second stage, edges between triangles are flipped if the angle of two vertices opposing the edge add up to more than $\pi$. For a curved surface, this operation can result in area, volume and shape changes. To limit this effect, flipping is only done for edges between neighboring triangles that make an angle $\theta < \pi/3$. Otherwise, the edge is split in the center where both triangles are subdivided into two triangles. In the last stage, edges between triangles that are too small ($S_A < S_{\max}$) are deleted, removing both triangles. This is repeated every $2\eta_c t_c / \sqrt{3}\gamma\Delta t$ timesteps. With remeshing, we assume that the nodes do not carry local information, but are merely considered as discretization points to represent a continuous surface.

### References


[1] Yuansheng Cao, Richa Karmakar, Elisabeth Ghabache, Edgar Gutierrez, Yanxiang Zhao, Alex Groisman, Herbert Levine, Brian A. Camley, and Wouter-Jan Rappel. Cell motility dependence on adhesive wetting. *Soft Matter*, 15:2043–2050, 2019.

[2] Françoise Brochard-Wyart and Pierre-Gilles de Gennes. Unbinding of adhesive vesicles. *Comptes Rendus Physique*, 4(2):281–287, 2003.

[3] Maxim Cuvelier, Jef Vangheel, Wim Thiels, Herman Ramon, Rob Jelier, and Bart Smeets. Stability of asymmetric cell division: A deformable cell model of cytokinesis applied to c. elegans. *Biophysical Journal*, 122(10):1858–1867, 2023.

[4] Steven Ongenae, Hanna Svitina, Tom E. R. Belpaire, Jef Vangheel, Tobie Martens, Pieter Vanden Berghe, Ioannis Papantoniou, and Bart Smeets. Active foam dynamics of tissue spheroid fusion. *bioRxiv*, 2024.

[5] Jean-Léon Maître, Hélène Berthoumieux, Simon Frederik Gabriel Krens, Guillaume Salbreux, Frank Jülicher, Ewa Paluch, and Carl-Philipp Heisenberg. Adhesion functions in cell sorting by mechanically coupling the cortices of adhering cells. *Science*, 338(6104):253–256, 2012.

[6] D. Fedosov, B. Caswell, and G. Karniadakis. Systematic coarse-graining of spectrin-level red blood cell models. *Computater Methods Applied Mechanics and Engineering*, 199:29–32, 2010.

[7] Bart Smeets, Tim Odenthal, Simon Vanmaercke, and Herman Ramon. Polygon-based contact description for modeling arbitrary polyhedra in the discrete element method. *Computer Methods in Applied Mechanics and Engineering*, 290:277–289, 2015.

[8] Kenneth A. Brakke. The surface evolver. *Experimental Mathematics*, 1:141–165, 1992.
